# Structure-aware deep learning predicts influenza antigenicity and guides vaccine strain recommendation

**DOI:** 10.64898/2026.08.02.742372

**Authors:** Xingyi Li, Chunyan Zhou, Kexin Xiao, Jialuo Xu, Xiangting Jia, Dongmin Zhao, Lei Chen, Yue Li, Jiajie Peng, Junnan Zhu, Yan Liu, Xuequn Shang, Huihui Kong

## Abstract

The continuous accumulation of genetic mutations in influenza A viruses (IAVs) drives antigenic drift, necessitating precise antigenic prediction for optimal vaccine strain selection. While sequence-based methods have advanced antigenic surveillance, they neglect the three-dimensional structural context that fundamentally dictates viral antigenicity. Here, we introduce Vir3D, which leverages ESMFold-derived structural information from amino acid sequences to precisely predict viral antigenicity and guide vaccine strain selection. Across both human H3 and highly pathogenic avian H5 subtypes, Vir3D not only accurately discriminates antigenic variants and infers pairwise antigenic distances, but also mechanistically delineates key structural residues driving viral immune evasion. In a decade-long retrospective analysis, Vir3D-prioritized vaccines consistently achieve broader antigenic coverage of circulating strains than World Health Organization (WHO) recommendations. Crucially, Vir3D successfully predicts that the emerging U.S. dairy cattle H5N1 virus (TX/24) remains antigenically stable relative to clade 2.3.4.4b vaccine strains, and subsequent wet-laboratory validation of hemagglutination inhibition (HI) assays definitively corroborates this finding. Overall, Vir3D establishes a powerful, structure-driven framework for proactive influenza surveillance and pandemic preparedness.

## Introduction

Influenza A viruses (IAVs) impose a substantial morbidity and mortality burden worldwide [1]. Human seasonal influenza is primarily dominated by the H3 subtype, which exhibits rapid antigenic drift and often evades vaccine-induced immunity, leading to recurring vaccine mismatches and severe seasonal epidemics [2]. Beyond human seasonal influenza, avian influenza viruses, particularly the H5 subtype, present a critical pandemic threat, causing widespread disruption to poultry and dairy production and endangering public health [3].

Effective control of IAVs relies fundamentally on timely vaccination, where protective efficacy is strictly dictated by the precise antigenic match of vaccine candidates with circulating viruses [4]. To guide vaccine strain selection, the World Health Organization (WHO) orchestrates the Global Influenza Surveillance and Response System (GISRS) to monitor viral evolution and antigenic drift worldwide [5]. Despite these extensive surveillance efforts, effective vaccine recommendation remains a formidable challenge, highlighted by historical vaccine mismatches [6–10]. To overcome rapid viral evolution and facilitate precise vaccine updates, there is an urgent demand for proactive antigenic prediction.

Although hemagglutination inhibition (HI) assays serve as the gold standard for evaluating the antigenicity of circulating isolates, they are fundamentally labor-intensive, time-consuming, and unsuited for high-throughput screening. Since antigenic drift is predominantly driven by mutations within the hemagglutinin subunit 1 (HA1) protein, the massive accumulation of high-throughput sequencing data naturally prompts the emergence of sequence-based antigenic predictors. Several studies have focused on sequence-based matrix completion to predict antigenic distances between viruses from partially observed HI matrices [11]. These methods incorporate sequence-derived information as prior constraints or regularization terms to infer missing entries. However, these approaches are fundamentally limited by their reliance on pre-existing HI measurements, making them inapplicable to target viruses without serological data. To overcome this limitation, recent studies have been developed for antigenic prediction directly from viral sequences, such as PREDAC [12], MFPAD [13], CNN-PSO [14], PREDAC-CNN [15], and AdaBoost [16]. These methods typically encode viral sequences using the physicochemical properties in AAindex and then utilize machine learning models for antigenic prediction.

Beyond the primary sequence, the three-dimensional (3D) structure of the HA1 protein dictates the biological function and ultimate antigenicity of IAVs [17, 18]. By fundamentally overlooking essential protein structural data, sequence-based models exhibit a strictly limited capacity for accurate antigenic prediction. This reliance on one-dimensional sequence data stems primarily from the extreme scarcity of experimentally resolved 3D structures for viral proteins. Fortunately, the rapid advancement of protein large language models has effectively overcome this critical bottleneck. By efficiently predicting high-fidelity 3D structures directly from primary sequences, these foundation models unlock unprecedented opportunities to develop structure-guided antigenic predictors.

In this study, we introduce Vir3D, a deep learning-based framework that leverages ESMFold-predicted protein structures to enable precise antigenic prediction and inform optimal vaccine strain selection. Specifically, Vir3D first translates the raw protein sequence of each virus into 3D struc-ture using ESMFold to construct protein graph. This graph is then processed through a geometric graph learning module to generate structural embeddings of each residue, intrinsically incorporating important site information related to antigenic characteristics to improve predictive accuracy. The proposed framework presents three key advantages. First, to the best of our knowledge, Vir3D is a pioneering framework for structure-aware antigenic prediction of IAVs, successfully harnessing the crucial features inherent to protein structures. Second, Vir3D is biologically informed, integrating important sites known to affect the antigenic properties of IAVs. Third, the geometric graph learning architecture natively captures complex spatial adjacency and geometric features of residues, while the explainable attention mechanism explicitly delineates the critical structural determinants driving immune escape, thereby providing mechanistic insights into viral evolution. Comprehensive evaluations demonstrate that Vir3D significantly surpasses existing sequence-based methods in antigenic predictive accuracy, while pinpointing the critical structural sites driving antigenic changes. Moreover, a decade-long retrospective study validates that Vir3D-selected vaccine candidates achieve superior antigenic coverage rates compared to WHO recommendations, establishing Vir3D as a vital decision-support framework for vaccine selection and pandemic surveillance. Most notably, Vir3D successfully predicts that the emerging U.S. dairy cattle H5N1 virus (TX/24) remains antigenically stable relative to the clade 2.3.4.4b vaccine strains. Subsequent wet-laboratory validation of HI assays has definitively confirmed that neither TX/24 nor its variants have undergone detectable antigenic drift. This not only supports the Re14 2.3.4.4b as an effective vaccine strain but also demonstrates Vir3D as a highly reliable computational framework for the rapid assessment of emerging viral threats.

## Results

### Overview of Vir3D

Vir3D is a structure-aware deep learning framework that enables accurate antigenic prediction of IAVs through geometric graph learning based on ESMFold-predicted protein structures (Figure 1). Given a viral HA1 sequence, its 3D structure is first predicted using the ESMFold protein language model and used to construct a protein graph, where residues are represented as nodes and inter-residue spatial proximities as edges. Node features encode intra-residue distances, directions, and angles, whereas edge features capture inter-residue distances, directions, and relative orientations. The constructed protein graph is then processed by a geometric graph learning module to generate structural embeddings of residues. Furthermore, these structural embeddings are functionally integrated with important site information, including glycosylation, receptor-binding, and antigenic epitope sites, before being aggregated into a global protein embedding via a multilayer perceptron (MLP). To evaluate antigenic relationships, the independent protein embeddings of both the target antigen and the reference antiserum are extracted through this pipeline. These paired embeddings are then concatenated, processed by a Transformer encoder layer, and fed into a terminal MLP for final antigenic prediction.

**Figure 1:**
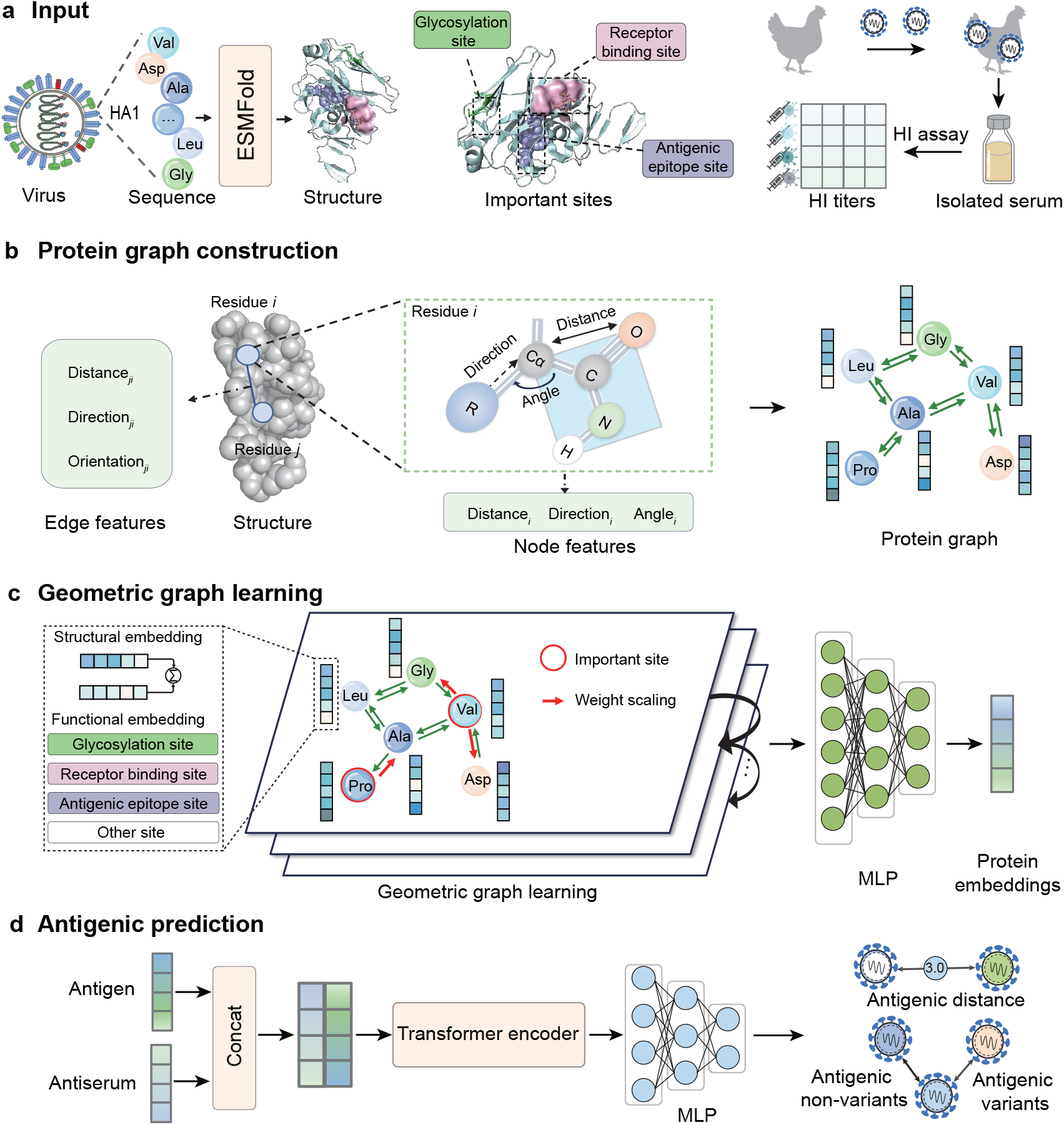
Overview of Vir3D. a) Vir3D takes three inputs: ESMFold-predicted protein structures derived from HA1 sequences, functionally important site information, and HI titers. b) For a given virus, the predicted structure is first used to construct a protein graph, in which individual residues are represented as nodes, inter-residue spatial proximities are encoded as edges, and geometric features are extracted directly from the protein structure. c) This graph is then processed through a geometric graph learning module to generate residue-level structural embeddings. To incorporate biologically meaningful context, these structural embeddings are integrated with functional embeddings derived from the important site information, and subsequently aggregated into a global protein embedding via an MLP. This pipeline yields an independent protein embedding for each virus. d) The independent embeddings of the target antigen and antiserum are subsequently concatenated, refined through a Transformer encoder layer, and fed into a final MLP for antigenic prediction.

### Vir3D robustly predicts the antigenic characteristics of influenza A viruses in cross-validation

To comprehensively evaluate predictive performance, we quantify the accuracy of Vir3D in inferring antigenic distances between virus pairs using Mean Absolute Error (MAE) and Root Mean Squared Error (RMSE). Simultaneously, we assess its capacity to discriminate antigenic variants from non-variants using the Area Under the Receiver Operating Characteristic Curve (AUC) and Area Under the Precision-Recall Curve (AUPRC) (see Methods for details). To ensure robust validation, we employ a five-fold cross-validation strategy with five independent repetitions (Methods and Figure S1). Additionally, we systematically benchmark Vir3D against a panel of competitors. This includes five sequence-based antigenic prediction models, namely MFPAD [13], AdaBoost [16], PREDAC [12], CNN-PSO [14], and PREDAC-CNN [15], alongside three baseline models encompassing SVM, CNN, and GCN (implementation details are described in Methods). Notably, Vir3D yields highly precise antigenic distance estimations (Figure 2a, MAE=0.715 and RMSE=0.941 for the H3 subtype, MAE=0.907 and RMSE=1.276 for the H5 subtype) and exhibits an exceptional capacity to identify antigenic variants (Figure 2b, AUC=0.887 and AUPRC=0.932 for the H3 subtype, AUC=0.925 and AUPRC=0.987 for the H5 subtype). These results confirm that the structure-aware representations learned via the geometric graph learning fundamentally drive superior predictions of antigenic characteristics.

**Figure 2:**
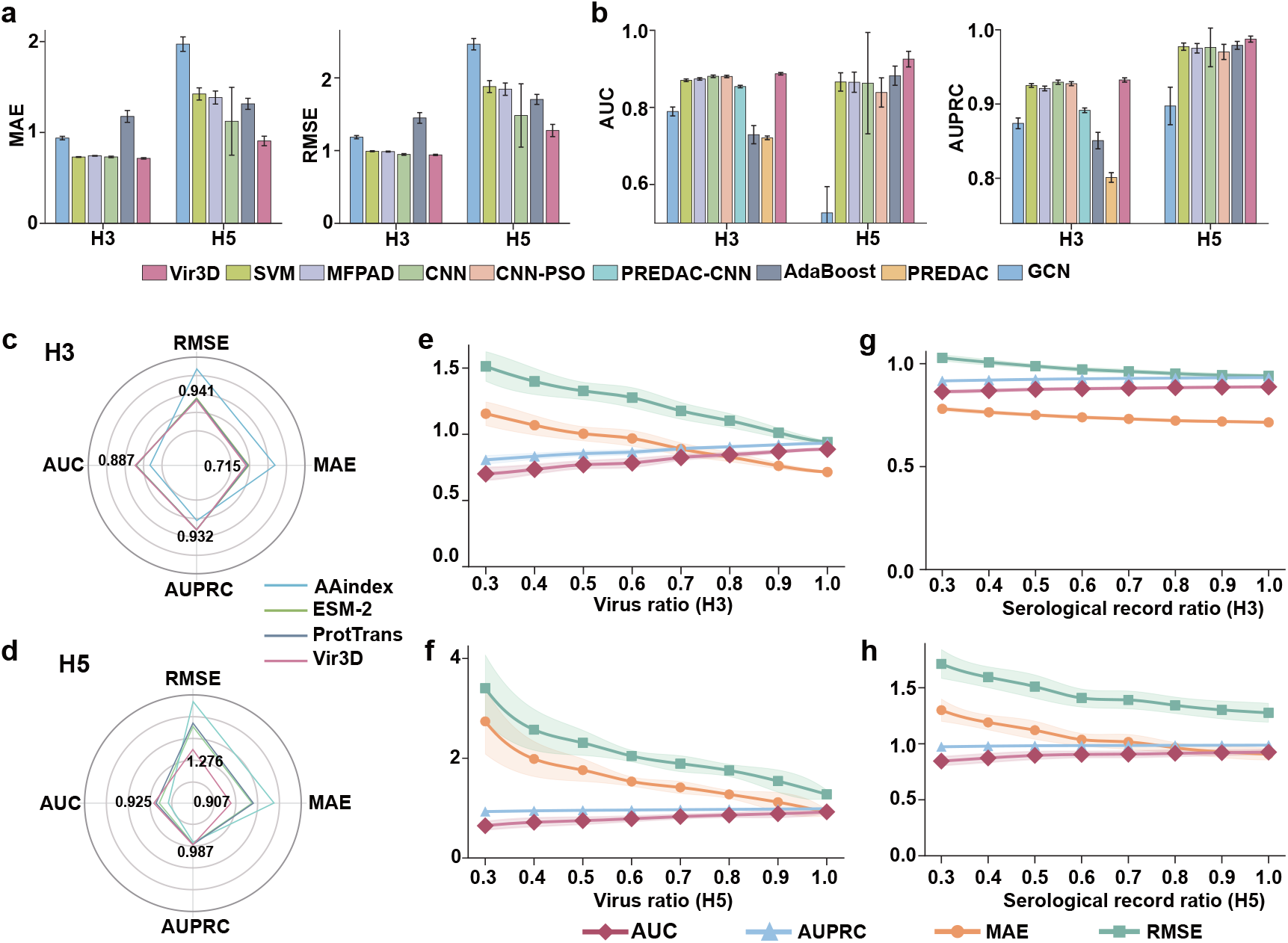
Performance evaluation for cross-validation. a) Comparative performance of Vir3D against five sequence-based methods (MFPAD, AdaBoost, PREDAC, CNN-PSO, and PREDAC-CNN) and three baselines (SVM, CNN, and GCN) in predicting antigenic distances. b) Comparative performance of Vir3D against five sequence-based methods and three baselines in discriminating antigenic variants from non-variants. c) Comparative performance of Vir3D against three different sequence-encoding methods (AAindex, ESM-2, and ProtTrans) for the H3 subtype. d) Comparative performance of Vir3D against three different sequence-encoding methods for the H5 subtype. e) Robustness of Vir3D evaluated by training on data subsets containing 30%–90% of the viruses, selected at random for the H3 subtype. f) Robustness of Vir3D evaluated by training on data subsets containing 30%–90% of the viruses, selected at random for the H5 subtype. g) Robustness of Vir3D evaluated by training on data subsets containing 30%–100% of the serological records, selected at random for the H3 subtype. h) Robustness of Vir3D evaluated by training on data subsets containing 30%–100% of the serological records, selected at random for the H5 subtype.

To further assess the critical contribution of structural features in characterizing antigenicity, we benchmark Vir3D against three prominent sequence-encoding baselines: AAindex [19], ESM-2 [20], and ProtTrans [21] (see Methods for details). Across both the H3 and H5 subtypes, Vir3D demonstrates consistent superiority in predicting antigenic distances and discriminating antigenic variants from non-variants (Figure 2c, d). Protein language models (ESM-2 and ProtTrans) show moderate effectiveness, whereas the traditional AAindex exhibits the poorest predictive performance. These findings definitively demonstrate that explicitly modeling 3D structural features provides a substantial accuracy advantage over sequence-only representations.

Furthermore, we validate the robustness of Vir3D under data scarcity using virus-centric and serological record-centric sampling strategies with retention rates varying from 30% to 100%. To simulate the practical absence of specific viral strains in surveillance, the virus-centric setting preserves serological records only when both the target antigen and the reference antiserum are included in the selected subset. Meanwhile, the serological record-centric setting randomly selects a subset of the available serological records, reflecting the natural incompleteness of serological measurements. Notably, Vir3D exhibits remarkable robustness, preserving high predictive accuracy despite a substantial reduction in training data, thereby underscoring its stability under data-limited conditions (Figure 2e–h).

### Vir3D accurately predicts antigenic characteristics of influenza A viruses in upcoming years

The timely identification of emerging antigenic variants and the precise characterization of circulating viral strains are paramount for global pandemic preparedness. To this end, the WHO has established National Influenza Centers worldwide to monitor antigenic evolution [5]. Accordingly, we implement a retrospective testing strategy designed to mimic real-world scenarios and predict the antigenic characteristics of circulating viruses in upcoming years (Methods and Figure S2). Under this framework, Vir3D demonstrates superior performance over competing methods across both the H3 and H5 subtypes in predicting antigenic distances (Figure 3a and b, MAE=0.836 and RMSE=1.074 for the H3 subtype, MAE=1.983 and RMSE=2.380 for the H5 subtype) and discriminating antigenic variants from non-variants (Figure 3c and d, AUC=0.820 and AUPRC=0.891 for the H3 subtype, AUC=0.739 and AUPRC=0.982 for the H5 subtype).

**Figure 3:**
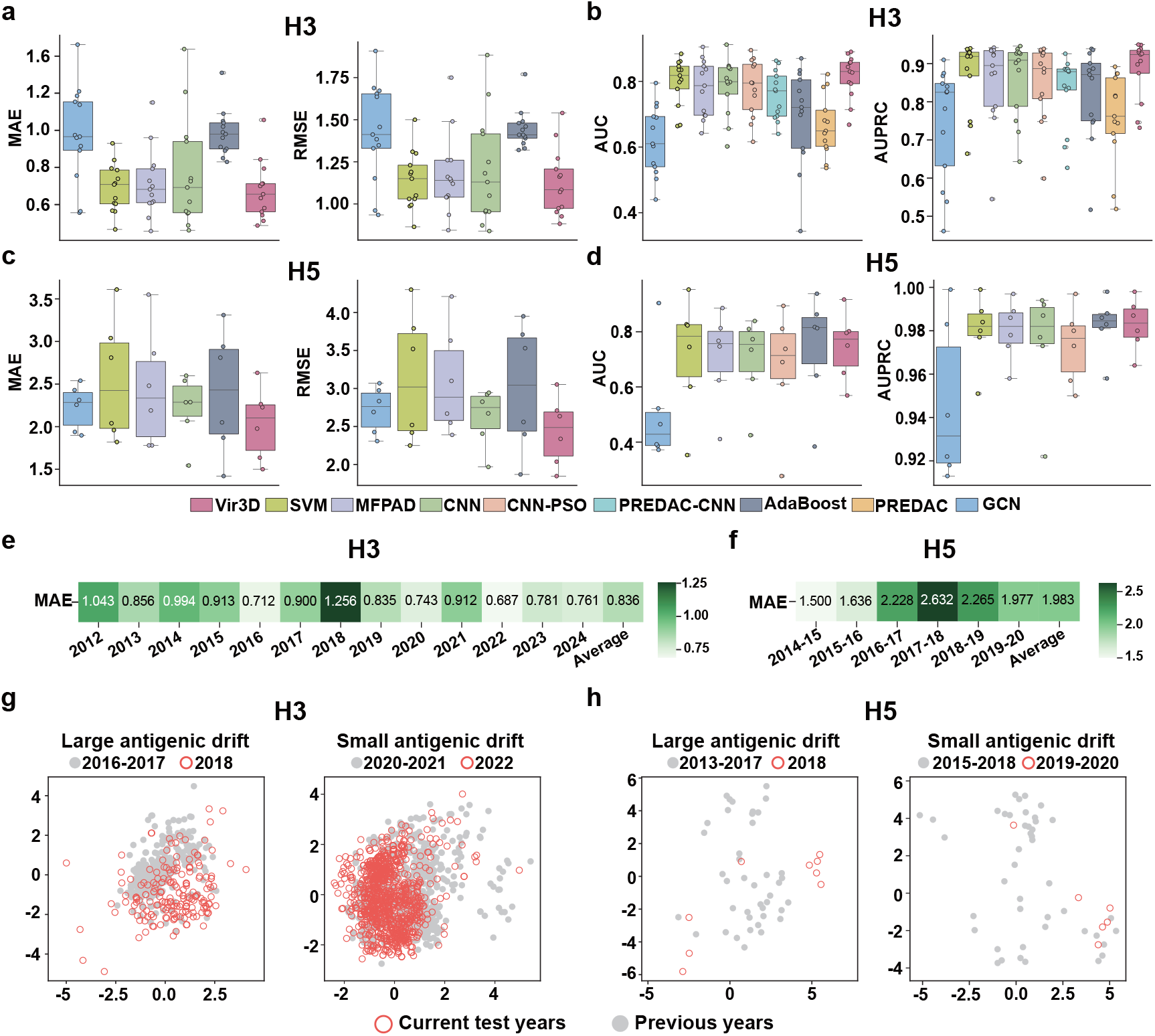
Performance evaluation for retrospective testing. a) Comparative performance of Vir3D against competing methods in predicting antigenic distances for the H3 subtype. b) Comparative performance of Vir3D against competing methods in predicting antigenic distances for the H5 subtype. c) Comparative performance of Vir3D against competing methods in discriminating antigenic variants from non-variants for the H3 subtype. d) Comparative performance of Vir3D against competing methods in discriminating antigenic variants from non-variants for the H5 subtype. e) MAE of Vir3D evaluated over 13 years for the H3 subtype. f) MAE of Vir3D evaluated over 7 years for the H5 subtype. g) Antigenic maps visualizing the antigenic drift of circulating viruses relative to viruses collected from the previous years for the H3 subtype using FluNexus. h) Antigenic maps visualizing the antigenic drift of circulating viruses relative to viruses collected from the previous years for the H5 subtype using FluNexus. For each subtype, the left map illustrates an instance of large antigenic drift, while the right map depicts a small antigenic drift.

While Vir3D generally demonstrates robust performance, it exhibits a noticeably elevated prediction error for both the H3 and H5 subtypes in 2018, excluding the initial test year with limited training data (Figure 3e, f). To investigate this anomaly, we employ Racmacs [22] and FluNexus [**li2026flunexus**] to visualize antigenic evolution trends, generating antigenic maps that depict the antigenic drift of circulating viruses relative to historical strains for the H3 and H5 subtypes (Figure 3g, h, and Figure S3). The resulting antigenic maps reveal that the 2018 circulating viruses (red circles) diverge significantly from those of earlier years (gray points). This substantial antigenic shift provides a biological explanation for the temporary decline in predictive accuracy. Although predictive accuracy temporarily declines during years with pronounced antigenic shifts, Vir3D remains robust across the remaining evaluation years and is most consistent during periods of limited antigenic drift (Figure 3e–h and Figure S3).

### Vir3D identifies significant sites associated with key antigenic regions of HA1

Accumulating evidence demonstrates that site-specific substitutions within HA1 sequences are major drivers of influenza antigenic evolution [23, 24]. Through the attention mechanism in the geometric graph learning framework (see Methods for details), Vir3D prioritizes sites most strongly associated with antigenic drift, thereby providing deep mechanistic insights into viral antigenic evolution.

For the H3 subtype, 27 of the top 30 critical sites prioritized by Vir3D map directly to key antigenic regions, encompassing epitopes A–E [23, 25] and receptor-binding regions (the 190-helix, 130-loop, and 220-loop) [26] (Figure 4a, c and g). These regions are widely recognized as primary determinants of IAV antigenicity, where substitutions have a great effect on the antigenic evolution of H3 and H5 viruses [22–24, 27–31]. Additionally, Vir3D identifies three high-ranking sites (sites 39, 282, and 309) residing outside these canonical regions. Structural analysis demonstrates that two of these sites are in close proximity to known epitopes in 3D space (with distances between carbon-alpha atoms *<* 7 Å), with site 282 positioning adjacent to epitope C, and site 309 in close proximity to epitope A (Figure 4g). Similarly, in the H5 subtype, 27 of the top 30 sites localize within key antigenic regions (Figure 4b, d and h), and two non-canonical sites exhibit structural proximity to established epitopes (site 109 to epitope E, and site 224 to epitope D). Overall, these findings validate Vir3D’s robust capability to capture and refine prior biological knowledge of critical sites through spatial contexts.

**Figure 4:**
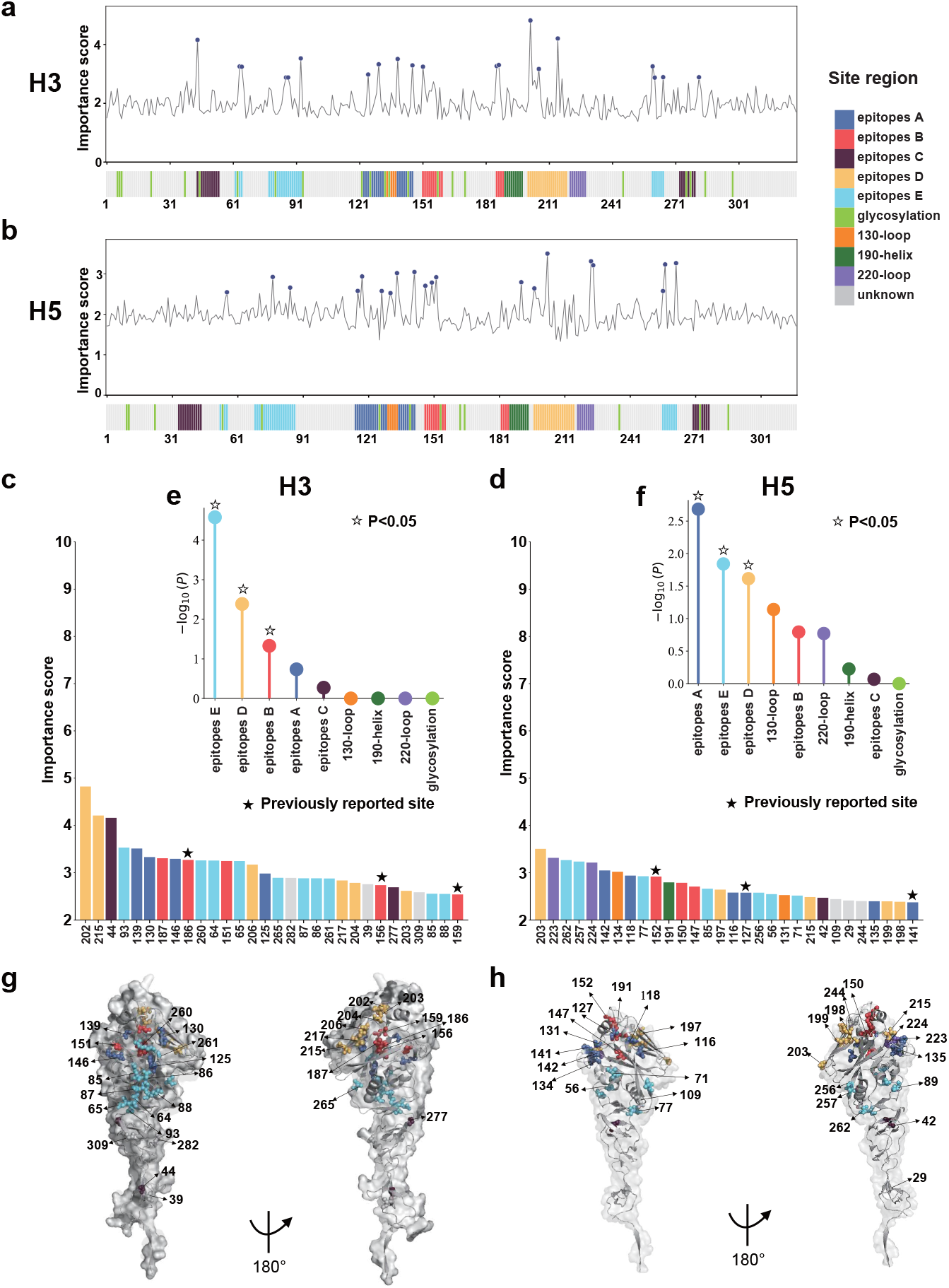
Important sites identified using Vir3D. a) Line plots comparing importance scores across HA1 amino acid sites for the H3 subtype. The top 30 sites exhibiting the highest importance scores are highlighted as blue dots. Sites not located within any key antigenic regions are categorized as ‘unknown’. b) Line plots comparing importance scores across HA1 amino acid sites for the H5 subtype. c) The top 30 sites with the highest importance scores for the H3 subtype. d) The top 30 sites with the highest importance scores for the H5 subtype. e) Enrichment analysis of the top 30 identified sites within specific antigenic regions for the H3 subtype. The statistical significance is quantified by the negative logarithm of the P-value. f) Enrichment analysis of the top 30 identified sites within specific antigenic regions for the H5 subtype. g) Structural mapping of the top 30 sites onto the HA1 protein for the H3 subtype (Protein Data Bank ID: 4KVN; A/Perth/16/2009). The top 30 sites are color-coded based on their corresponding antigenic regions. h) Structural mapping of the top 30 sites onto the HA1 protein for the H5 subtype (Protein Data Bank ID: 4K62; A/Indonesia/5/2005).

To further investigate the biological relevance of these top-ranking sites in antigenic evolution,we perform a detailed analysis of representative sites. For the H3 subtype, sites 156, 159, and 186 within epitope B exhibit high importance scores (Figure 4a, c). Existing evidence confirms that sites 156 and 159 exert a strong impact on antigenic evolution [24], with the Y159F substitution has been identified as a major driver of antigenic drift from the FU02 to CA04 antigenic clusters [32]. Additionally, the G186D mutation is identified as a key determinant of antigenic drift by stabilizing the N190 side-chain conformation through a critical hydrogen bond, which facilitates a direct interaction between hemagglutinin and human receptors and thereby drives the evolution of the receptor-binding mode [33]. Regarding the H5 subtype, Vir3D assigns notably high attention weights to site 152 within epitope B, as well as sites 141 and 127 within epitope A (Figure 4b, d). Experimental evidence confirms that site 141 serves as a major antigenic determinant for the 2.3.4.4 clade of the H5 subtype [28, 30]. Meanwhile, antigenic evolution studies of H5 avian influenza viruses consistently identify immune escape mutants harboring substitutions at site 152, highlighting its pivotal role in mediating immune evasion and driving antigenic drift [28]. Moreover, site 127 within epitope A is shown to be a critical determinant of antigenic escape for the H5 subtype [27].

Furthermore, statistical analysis reveals that the critical sites predicted for the H3 subtype are significantly enriched within epitopes B, D, and E (Figure 4e), whereas those for the H5 subtype are predominantly enriched in epitopes A, D, and E (*P <* 0.05) (Figure 4f). Biologically, epitope B has been established to exhibit the highest immunodominance for the H3 subtype [24, 25], while epitope A serves as the primary hotspot for antigenic variation in H5 viruses and is predominantly targeted by the host immune response [29–31]. Consequently, these results demonstrate the capacity of Vir3D to accurately delineate the subtype-specific, immunodominant epitopes that drive antigenic evolution.

### Vir3D enables optimal vaccine recommendation based on coverage rates

The ultimate goal of antigenic prediction is to enable precise vaccine strain recommendation, thereby empowering effective epidemic prevention and control. Fundamentally, vaccine efficacy is dictated by the antigenic match between the vaccine strain and circulating viruses. To evaluate the utility of Vir3D in prospective vaccine strain selection, we design a retrospective evaluation that faithfully mirrors the decision-making process of the WHO Vaccine Composition Meetings: given surveillance data available up to the time of selection, the optimal candidate strain is defined as that which would confer the broadest cross-protective coverage against viruses ultimately circulating in the forthcoming year. Since future epidemic strains are inherently unknown at the time of selection but typically evolve from currently dominant antigenic clusters, targeting the antigenic centroid of recently circulating viruses is a rational strategy to maximize broad cross-protective immunity against forthcoming variants. Accordingly, we introduce the coverage rate (see Methods for details) as a quantitative metric to assess the proportion of recently circulating viruses antigenically matched to a given candidate strain, thereby validating the prophylactic efficacy of the recommended vaccine strain.

A fundamental obstacle to computing reliable coverage rates from raw surveillance data is the inherent sparsity of the global HI titer matrix: because most antigens lack direct serological mea-surements, data-driven coverage rate estimation from HI data alone is statistically unstable and effectively infeasible at scale. We address this limitation by leveraging Vir3D to predict pairwise antigenic distances across the full virus landscape, enabling the calculation of statistically robust coverage rates even in the absence of HI measurements. To verify that these computationally derived coverage rates faithfully reflect those derived from HI titers, we conduct a rigorous correlation analysis against coverage rates calculated exclusively from available HI titers (see Methods for details). The distribution of Vir3D-derived coverage rates closely mirrors that of HI-based coverage rates (Figure S4a), residuals are tightly centered around zero (Figure S4b), and both Pearson and Spearman correlation analyses confirm a strong positive association (Pearson correlation coefficient [34] *R* = 0.842, *P <* 0.01; Spearman’s rank correlation [35] *ρ* = 0.854, *P <* 0.01). Collectively, these results demonstrate that coverage rates derived from Vir3D-predicted antigenic distances serve as a reliable and accurate surrogate for those derived from HI measurements, and can thus be confidently employed to benchmark the cross-protective efficacy of recommended vaccine strains in subsequent evaluations.

To obtain Vir3D-based recommendations, optimal candidate strains are identified using coverage rates predicted by Vir3D trained exclusively on historical data available prior to the target year. The virus exhibiting the highest predicted coverage rate is then designated as the Vir3D-recommended vaccine strain for the upcoming year (Figure 5a). To rigorously evaluate the efficacy of these recommended strains, we benchmark them against the WHO-recommended strains for both the northern and southern hemispheres (representing the real-world baseline) and the Oracle-recommended strains (representing the theoretical upper bound of achievable performance). Specifically, the Oracle strategy selects the optimal vaccine strain based on coverage rates calculated by Vir3D trained on data inclusive of the target year, effectively representing a perfectly informed, retrospective selection. The specific strains recommended by each strategy in each evaluated year are detailed in Tables S1 and S2.

**Figure 5:**
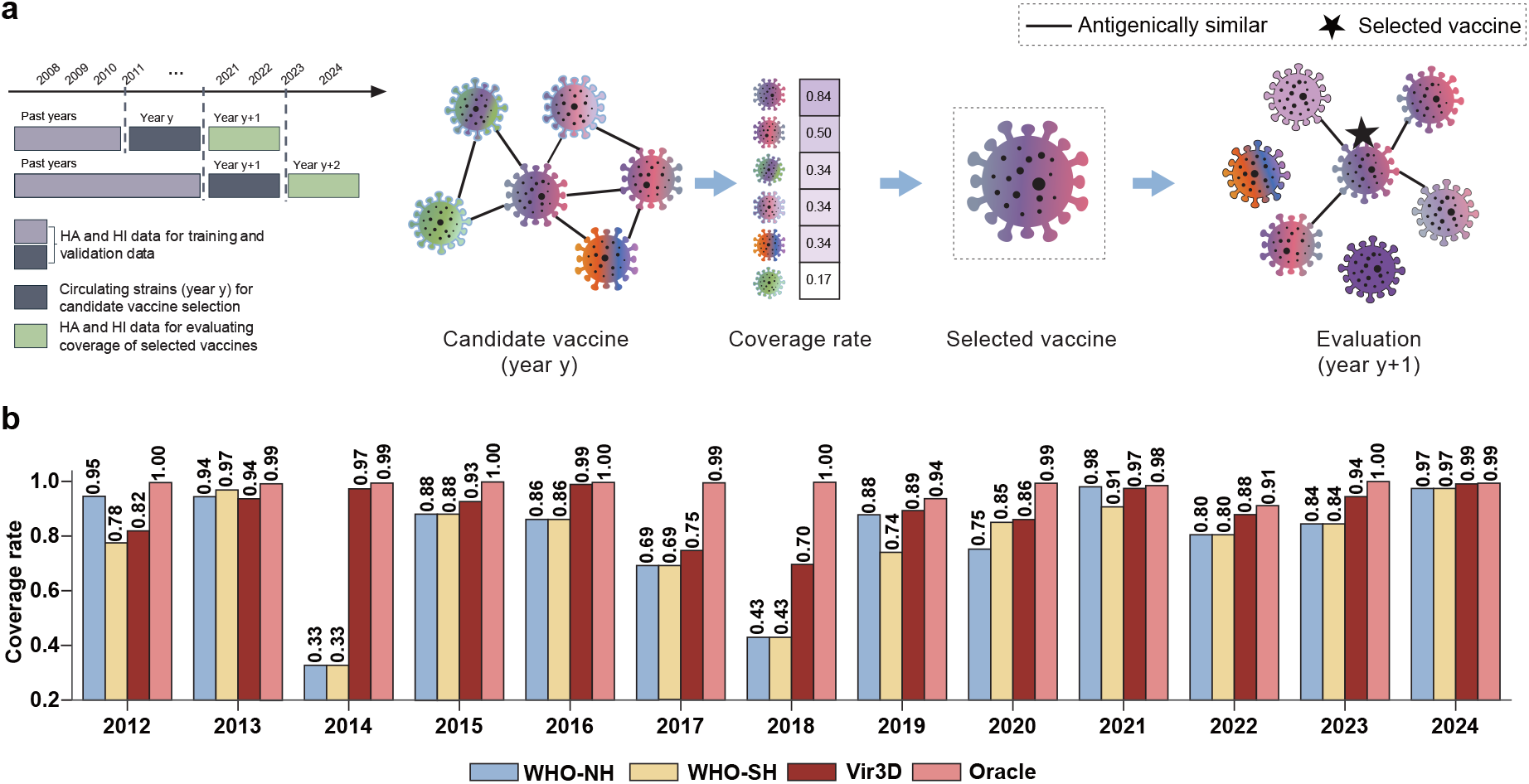
Vaccine recommendation based on coverage rates. a) Schematic of the vaccine recommendation workflow. Using only historical HA and HI data available before the target year *y* + 1, Vir3D identifies the candidate virus with the highest predicted coverage rate among strains circulating in year *y* as the recommended vaccine strain. The recommended strain is then evaluated against viruses circulating in the target year *y* + 1 using the coverage rate. b) Coverage rates of vaccine strains recommended by the WHO for the northern hemisphere (WHO NH) and southern hemisphere (WHO SH) influenza seasons, by Vir3D, and by the Oracle upper bound (2012–2024).

The evaluation results demonstrate the exceptional robustness and accuracy of Vir3D (Figure 5b). As expected, the vaccines recommended by the Oracle consistently achieve near-perfect coverage, confirming its role as a theoretical upper bound. Crucially, Vir3D-recommended strains outperform WHO recommendations for both the northern and southern hemispheres in 10 out of 13 evaluated years, and closely approach the theoretical upper bound established by the Oracle. The practical significance of Vir3D is most evident in the 2014 scenario. In this evaluation, the WHO-recommended A/Texas/50/2012 (H3N2) strain exhibits a severely compromised coverage rate of 0.33, which corroborates documented reports of severe antigenic mismatch during that year [6–10]. Conversely, the Vir3D-recommended vaccine strain attains a coverage rate of 0.97. Overall, these results validate the potential of Vir3D as a high-fidelity decision-support tool for prospective vaccine strain selection in real-world surveillance.

### Vir3D predicts antigenic stability of the dairy cattle H5 virus confirmed by wet-laboratory validation

Clade 2.3.4.4b H5 viruses are circulating globally and remain a pandemic concern. In March 2024, H5N1 was reported in dairy cattle in the United States (U.S.) and has continued to circulate. To assess whether the U.S. dairy cattle H5N1 virus exhibits antigenic divergence from 2.3.4.4b H5 viruses, we compare the first reported dairy cattle isolate, A/dairy cattle/Texas/24-008749-001-original/2024 (TX/24), with Re14 2.3.4.4b, a clade 2.3.4.4b vaccine strain widely used in China (Figure 6a). HA1 sequence comparison between TX/24 and Re14 2.3.4.4b identifies amino acid differences at sites 104, 115, 188, 195, and 210. In addition, analysis of all available U.S. dairy cattle H5N1 HA1 sequences in GISAID (as of March 5, 2026) indicates that the HA1 region remains largely conserved, with recurrent variation observed only at sites 88 and 131 (Figure S5).

**Figure 6:**
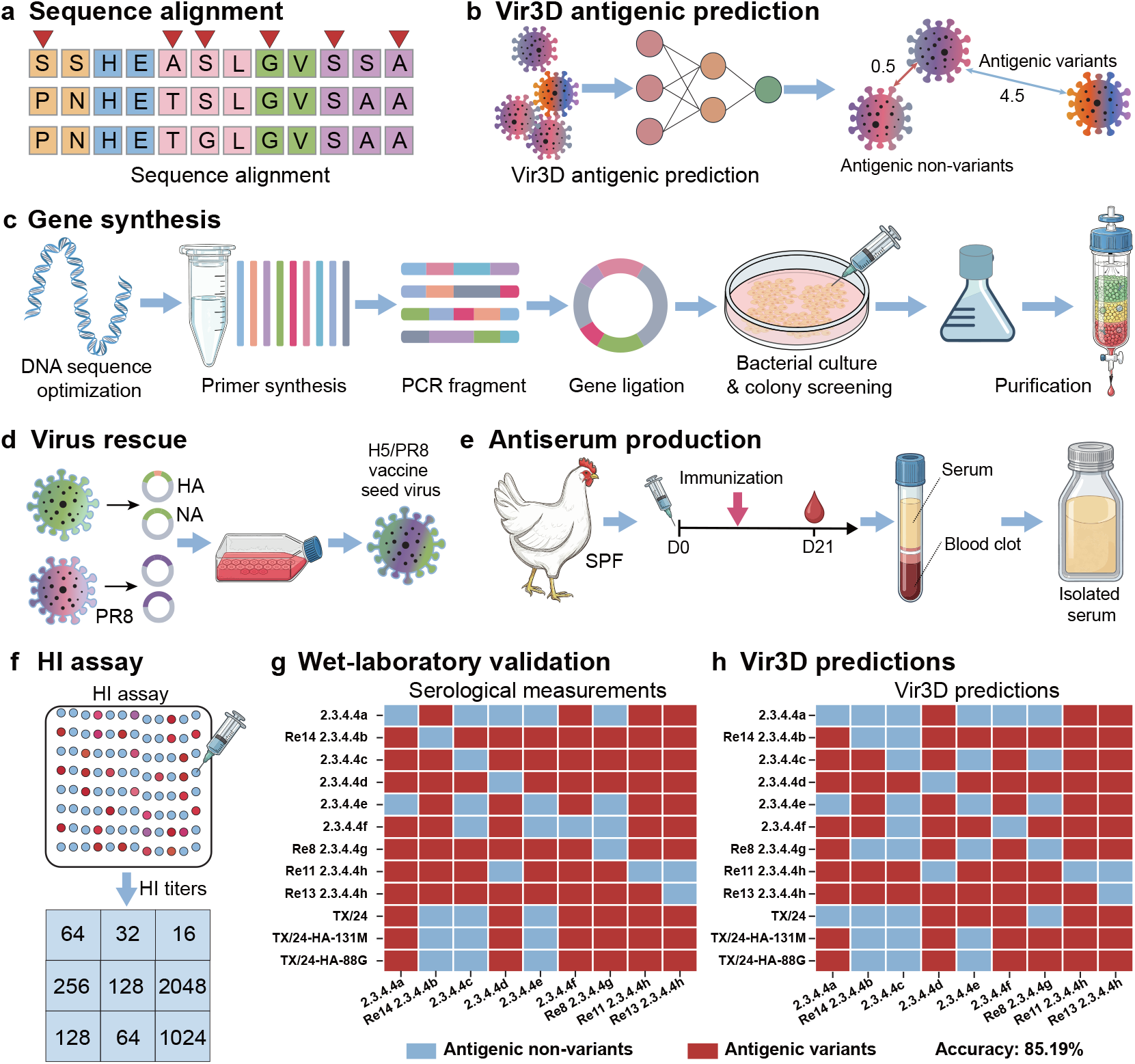
Workflow for antigenic evaluation of the U.S. dairy cattle H5N1 influenza virus. a) HA1 sequences are collected and aligned to identify amino acid differences. b) Vir3D is applied to predict antigenic distances between virus pairs based on HA1 structural features, and to determine whether viruses are classified as antigenic variants or antigenic non-variants. c) Genes of TX/24 and its mutants are synthesized and cloned into a reverse genetics vector. d) Reassortant viruses are rescued using a reverse genetics system containing the HA of interest, TX/24 neuraminidase (NA), and A/Puerto Rico/8/1934 (PR8) internal genes. e) Antisera are generated in specific-pathogen-free (SPF) chickens through immunization and serum collection. f) HI assays are performed to measure HI titers, calculate antigenic distances, and determine whether viruses are discriminated as antigenic variants or non-variants. g) Wet-laboratory validation using HI assays. The heatmap shows the results for discriminating antigenic variants from non-variants, derived from cross-reactive HI titers between representative H5 clade viruses and the TX/24 isolate along with its variants (TX/24-HA-88G and TX/24-HA-131M). h) Prediction of antigenic variants and non-variants using Vir3D. The heatmap shows the corresponding results inferred by Vir3D between representative H5 clade viruses and the TX/24 isolate and its variants.

Given these sequence differences, we utilize Vir3D to predict antigenic relationships among Re14 2.3.4.4b, TX/24, and TX/24 variants carrying the D88G (TX/24-HA-88G) or V131M (TX/24-HA-131M) (Figure 6b). Notably, the computational predictions demonstrate no meaningful antigenic shift among these viruses (Figure 6h, Figure S6a, and Table S3). To experimentally corroborate these computational findings, we synthesize the TX/24 HA, introduce the D88G or V131M substitution, rescue reassortant viruses carrying the indicated mutant HA, the TX/24 neuraminidase, and PR8 internal genes, and assess their antigenic properties using HI cross-reactivity assays with avian antisera (Figure 6c–f). Consistent with the Vir3D predictions, HI assays indicate no detectable antigenic differences among Re14 2.3.4.4b, TX/24, TX/24-HA-88G, and TX/24-HA-131M (Figure 6g, Figure S6b, and Table S4).

Taken together, these biological and computational findings confirm that the U.S. dairy cattle H5N1 virus remains antigenically stable despite the observed substitutions, and that the Re14 2.3.4.4b strain retains serological cross-reactivity against TX/24 and the tested variants. Beyond evaluating this specific pandemic preparedness concern, this case study underscores the power of Vir3D as a highly reliable framework for the rapid antigenic assessment of emerging viruses in real-world surveillance scenarios.

## Discussion

Antigenic prediction and vaccine recommendations play a crucial role in influenza control. Traditional serological assays, such as HI assays, have been widely used for antigenic evaluation due to their reliability. However, they are inherently labor-intensive, time-consuming, and low-throughput. These constraints have necessitated a paradigm shift toward predictive computational methods that can rapidly assess the antigenicity of IAVs. Antigenic drift, primarily driven by mutations in the HA1 protein, along with the rapid accumulation of high-throughput sequencing data, has paved the way for the development of sequence-based antigenic predictors. Despite these advances, sequence-based methods often fail to fully capture the virus’s complex antigenic characteristics. This is fundamentally because the 3D structure of the HA1 protein is critical for the biological function and ultimate antigenic characteristics of IAVs [17, 18].

Recognizing this structure-function relationship as central to viral immune evasion, we have developed Vir3D, a deep learning-based antigenic predictor that leverages ESMFold-predicted protein structures to enable precise antigenic prediction and inform optimal vaccine strain selection. Vir3D shifts the reliance from one-dimensional sequence analysis to a highly interpretable, three-dimensional structural perspective, and provides profound mechanistic insights into viral immune evasion. Comprehensive evaluations across both human seasonal H3 and highly pathogenic avian H5 subtypes reveal that Vir3D achieves highly accurate estimations of antigenic distances and exhibits a robust capacity to discriminate antigenic variants from non-variants. Critically, Vir3D maintains high predictive fidelity in prospective settings, accurately predicting the antigenic characteristics of viruses circulating in subsequent years, underscoring the effectiveness of structural features in modeling viral evolution. Beyond antigenic prediction, Vir3D provides mechanistic insights by pinpointing important sites within HA1 that have dominant impacts on antigenic changes. Furthermore, in a decade-long retrospective evaluation, Vir3D-recommended vaccine strains achieve superior antigenic coverage rates compared with WHO-recommended strains in 10 of 13 evaluated years, highlighting its potential for vaccine optimization and pandemic surveillance. Most importantly, the consistency between Vir3D predictions and the HI cross-reactivity assays confirms the antigenic stability of the circulating U.S. dairy cattle H5N1 virus. This not only supports the Re14 2.3.4.4b strain as an effective vaccine strain but also demonstrates Vir3D as a highly reliable framework for the rapid assessment of emerging viral threats.

Looking ahead, further advancements will depend on voluntary data sharing and closer integration with wet-laboratory validation, creating an iterative framework in which wet-laboratory findings refine model parameters and computational predictions guide subsequent experimental testing. Such a synergistic framework has the potential to further strengthen the role of Vir3D in real-time virological surveillance, vaccine updating, and preparedness for emerging influenza viruses.

## Materials and Methods

### Data collection and preprocessing

HA sequences of influenza H3 viruses are obtained from Smith et al. and the GISAID database [36]. For influenza H5 viruses, HA sequences are retrieved from the GISAID database [36] and supplemented with additional sequences derived from virus isolates collected in China and processed in our BSL-3 laboratory. To preprocess these sequences, sequences for each subtype are aligned separately using MAFFT, with A/Aichi/2/1968 (H3N2) (isolate ID: EPI_ISL_123225) and A/Vietnam/1203/2004 (H5N1) (isolate ID: EPI_ISL_10656749) used as reference strains for H3 and H5, respectively. After alignment, only the HA1 subunit is retained, and the signal peptides (16 and 16 amino acids for H3 and H5 viruses, respectively) are removed. In addition, sequences containing ambiguous amino acid codes (‘B’, ‘Z’, ‘J’, or ‘X’), more than 10% gaps, or terminal gaps at both ends are discarded. This standardized preprocessing pipeline is implemented in the FluNexus platform (https://flunexus.com/DataPreprocessing.html). As a result, the HA1 dataset for the H3 subtype comprises 149,383 sequences of 328 amino acids (1968–2025). Of these, 143,067 sequences lack paired serological data and are therefore restricted to the vaccine recommendation analysis, collectively constituting the candidate repertoire for strain screening. The final HA1 dataset for the H5 subtype comprises 4,107 sequences of 317 amino acids (1996–2025), of which 4,030 are HA1 sequences of U.S. dairy cattle H5N1 viruses.

HI titers for the H3 subtype are acquired from the Worldwide Influenza Centre (WIC) at the Francis Crick Institute [37] and Smith et al. [22]. For the H5 subtype, in-house HI titers are generated in our BSL-3 laboratory and are archived in the FluNexus repository [**li2026flunexus**] (https://github.com/xingyili/FluNexus-methodbox/tree/main/H5_data). Invalid HI titers denoted by ‘*<*’ are excluded, while titers indicated as ‘*>*512’ are treated as twice the value. To convert HI titer data into quantitative measurements, antigenic distances for antigen-antiserum pairs are calculated using the Normalized Antigenic Distance (NAD) as follows:

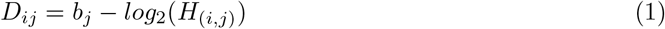

where *b*_*j*_ denotes the homologous titer of antiserum *j* and *H*_(*i,j*)_ represents the heterologous titer of antigen *i* against antiserum *j*. For each antigen-antiserum pair, the antigen represents a circulating virus isolate, while the antiserum is derived from the reference virus. Pairs lacking homologous titers or corresponding HA1 sequences are discarded. Valid pairs with an NAD distance ≥ 2 are classified as antigenically distinct [15, 22, 38] (data statistics are shown in Figure S7). Following refinement, we obtain 46,886 NAD values derived from 6,316 antigens and 214 antisera spanning from 1968 to 2024 for H3 subtype, and 1,848 NAD values corresponding to 77 antigens and 24 antisera spanning from 1996 to 2020 for the H5 subtype.

### Protein structure prediction model (ESMFold)

ESMFold is a protein language model designed to predict structures directly from protein sequences. Compared to other state-of-the-art models, ESMFold offers a favorable balance between computational efficiency, structural accuracy, and lightweight architecture, thereby enabling fast and scalable structure prediction across large-scale datasets. Leveraging these advantages, we employ ESMFold to efficiently generate HA1 protein structures, which serve as structural inputs for the geometric deep learning to elucidate the relationship between spatial conformation and antigenic characteristics.

### Protein graph construction

We model each protein structure as a graph G = (V, E), where nodes represent individual residues and edges encode spatial proximity relationships. Specifically, an edge connects nodes *i* and *j* if the Euclidean distance between their *C*_*α*_ atoms is less than a predefined threshold *d*_*c*_ (set to 8 Åfor H3 and 6 Åfor H5). The node and edge features in the graph are derived from a local coordinate system based on the backbone atoms of the residue. Given the coordinates of node *i*, 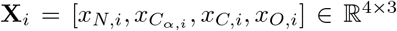, the backbone orientation vectors *u*_*i*_ and *v*_*i*_, and the local coordinate system **R**_*i*_ = [*b*_*i*_, *n*_*i*_, *t*_*i*_] are defined as:

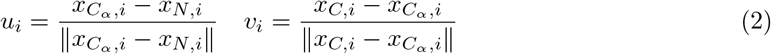

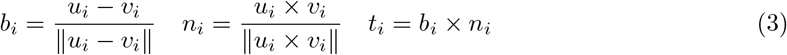

where *b*_*i*_, *n*_*i*_, and *t*_*i*_ denote the backbone direction, the normal vector, and the secondary tangent vector of the residue, respectively. Based on this local coordinate system, the node and edge features are defined as follows:

i. Node features. Node features integrate multiple geometrical features to comprehensively represent the structural characterization of each residue, encompassing angle, distance, and direction features. First, to capture backbone conformational geometry, dihedral angles (*ϕ*_*i*_, *ψ*_*i*_, *ω*_*i*_) and bond angles (*α*_*i*_, *β*_*i*_, *γ*_*i*_) are incorporated, with their sine and cosine values encoded as angle features. Second, to capture intra-residue spatial information, interatomic distances are encoded using radial basis functions (RBFs) as RBF(||*a*_*i*_ − *b*_*i*_||), where 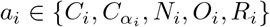 and 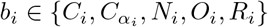 denote the atoms in residue *i* and *a* ≠*b*. Third, to characterize spatial orientation within the local coordinate system, the direction of each atom (*N, C, O, R*) relative to *C*_*α*_ is normalized and projected onto the local coordinate, serving as the direction feature 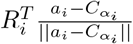.
ii. Edge features. Analogous to node features, edge features encode geometric relationships between two residues through orientation, distance, and direction features. For residues *i* and *j*, the relative orientation is represented as follows:

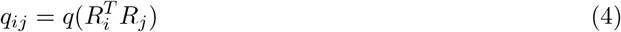

where *q*(·) denotes a quaternion function, mapping a rotation matrix to its quaternion representation [39]. Meanwhile, distance features are defined as RBF(∥*a*_*i*_ − *g*_*j*_∥) for 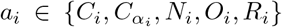 and 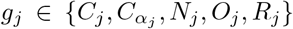, where *a*_*i*_ and *g*_*j*_ denote atoms in residues *i* and *j*, respectively.Additionally, direction features are defined as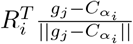, denoting the relative spatial positions of atoms in residue *j* in the local coordinate system of residue *i*.

### Geometric graph learning

To capture the spatial geometric characteristics of protein structures, the constructed protein graph is processed through graph neural network (GNN) layers of geometric graph learning. Meanwhile, to model multi-scale residue features, the attention modules are employed at different levels, including node, edge, and global context levels.

i. Node update. Given the demonstrated effectiveness of transformer architecture in modeling both sequence and graph data [40], we utilize its multi-head attention mechanisms to enable efficient message passing among nodes in the protein graph. At the *l*-th layer, the feature of node *i* and the edge feature from node *j* to *i* are denoted as 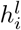and 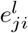, respectively, which are projected into a *d*-dimensional latent space prior to the GNN layer. Subsequently, the node features are updated through the following process:

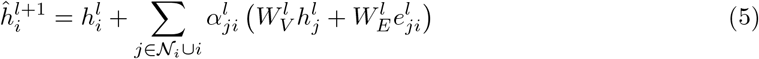

where *N*_*i*_ represents the neighbors of node *i* and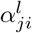 denotes the attention weight, formulated as follows:

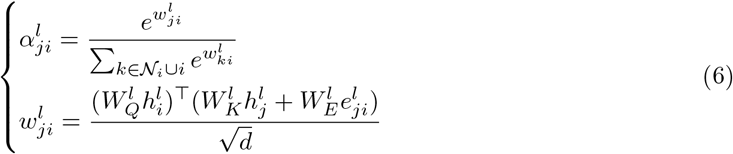

where 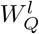, 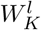, and 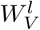 denote the learnable weight matrices that map node features into query, key, and value representations, respectively. 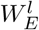 represents the learnable weight matrices designed to effectively incorporate the geometric information encoded in the edges into the message aggregation process. Multiple attention heads are applied in parallel, and their outputs are concatenated to form the updated node representation.
ii. Edge update. The edge features are updated using the information from the neighboring nodes.

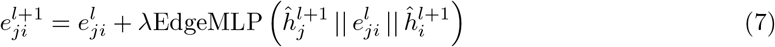

where *λ* ≥ 1 is a hyperparameter indicating the importance of the source node. To prioritize biologically significant sites, we assign *λ >* 1 specifically when the source node is located within an important region (glycosylation, receptor binding, and antigenic epitope sites), amplifying the contribution of the important sites during the update process. EdgeMLP represents the MLP operation used for edge update, and || denotes the concatenation operation.
iii. Global context attention. While node-level and edge-level features effectively capture local geometric relationships within neighborhood interactions, incorporating global structural information is essential for learning comprehensive residue representations. However, calculating global attention introduces significant computational complexity. To address this challenge, we implement an efficient global context module that first computes a global context vector, which is subsequently employed for node embedding with a gate attention [41]:

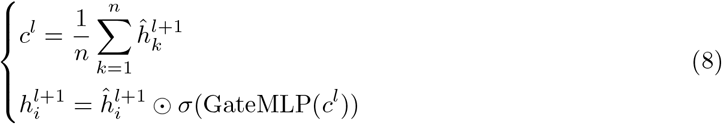

where *n* denotes the number of nodes in a protein graph, ⊙ indicates element-wise multiplication, *σ* is the sigmoid activation function, GateMLP represents a gating MLP for adaptively gating the global context, and 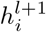 denotes the structural embedding of residue *i* generated by the *l*-th geometric graph learning layer, capturing the structural information.

To endow Vir3D with prior knowledge of critical functional sites, we construct learnable functional representations for each residue. Specifically, the biological attributes of residues are classified into 10 distinct categories, including glycosylation sites, receptor-binding sites (130-loop, 190-helix, and 220-loop), antigenic epitopes A–E, and non-specific background sites. Each category is mapped to a unique integer index and projected into a dense continuous space using a learnable lookup table. This yields a trainable functional embedding vector for each site, which is subsequently integrated with the structural embedding via element-wise addition. This design allows the model to dynamically learn the context-dependent influence of these critical antigenic regions during training. Subsequently, the resulting embeddings are processed through an MLP to derive a comprehensive, global protein embedding.

To predict the antigenic distances between antigens and antisera, the protein embeddings of the antigen and antiserum are concatenated and input into a transformer encoder layer to capture the non-linear association between them. Subsequently, the resulting embeddings are passed through an MLP for regression, enabling end-to-end prediction of antigenic distance for antigen-antiserum pairs. Moreover, antigen-antiserum pairs with a predicted NAD distance ≥ 2 are classified as antigenically distinct, whereas those with a predicted NAD distance *<* 2 are classified as antigenically similar.

### Implementation and evaluation

Vir3D is implemented using the PyTorch 1.13.1 framework, which incorporates a geometric graph network consisting of a 2-layer GNN with 64 hidden units. The attention mechanism within the geometric graph network employs multi-head attention with 4 attention heads. For model optimization, we utilize the AdamW optimizer. Meanwhile, the training process is constrained to a maximum of 80 epochs, with a dropout rate of 0.2 applied to mitigate overfitting. Notably, to facilitate model optimization and mitigate issues arising from the large magnitude of raw antigenic distances, we train and test Vir3D using standardized values. Specifically, the raw values are scaled using a standardized transformation with predefined parameters: *µ* = 0 and *σ* = 1 for the H3 subtype, and *µ* = 2 and *σ* = 2 for the H5 subtype.

To systematically assess the performance of Vir3D, we employ a comprehensive evaluation strategy comprising five-fold cross-validation and retrospective testing (Figs. S1 and S2). In the five-fold cross-validation, the dataset is randomly partitioned into five subsets. In each iteration, the model is trained on four folds and tested on the remaining fold. This operation is repeated five times to ensure robust performance assessment. To evaluate the real-world applicability of Vir3D under prospective surveillance conditions, we conduct retrospective testing. For each target year, Vir3D is trained exclusively on HI titer and HA1 sequence data available prior to that year, then tested on circulating isolates of the target year. Given the scarcity of historical data in earlier years (Figs. S8 and S9), we select 13 test years for H3 (2012–2024) and 7 test years for H5 (2014–2020) subtype evaluations. Notably, due to the relatively slow antigenic evolution of H5 viruses [31] and associated data sparsity, each testing round for H5 subtype aggregates two consecutive years. For each test round, the model is trained on the antigen-antiserum pairs spanning from the earliest available year through year *y* − 1, and subsequently tested on pairs of H3 subtype from year *y* or pairs of H5 subtype from years *y* and *y* + 1.

Meanwhile, to compare the performance of Vir3D to other methods under the cross-validation and retrospective testing strategies, we compute four key metrics: MAE and RMSE for evaluating the ability to predict antigenic distances, and AUC and AUPRC for assessing the performance to discriminate antigenic variants from non-variants. All metrics are first calculated for each individual fold or test year and subsequently aggregated via weighted averaging to evaluate the overall performance.

For the experiment of antigenic distance prediction, MAE_*k*_ and RMSE_*k*_ for fold *k* or test year *k* are calculated as follows:

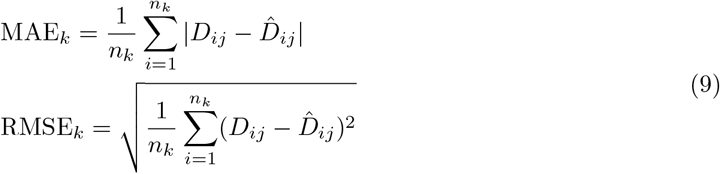

where *n*_*k*_ is the total number of antigen-antiserum pairs in test fold *k* or year *k, D*_*ij*_ represents the antigenic distance between antigen *i* and antiserum *j* derived from HI titers, and 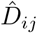 denotes the corresponding antigenic distance predicted by the model.

For the binary classification task of discriminating antigenic variants from non-variants, each antigen-antiserum pair is classified as either antigenically distinct (positive sample) or antigenically similar (negative sample). The AUC_*k*_ for fold *k* or test year *k* is calculated as the area under the Receiver Operating Characteristic curve, which evaluates the True Positive Rate (TPR) versus the False Positive Rate (FPR) across all possible classification thresholds. The TPR and FPR are defined as:

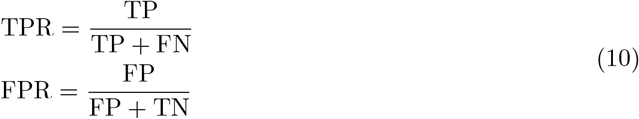

where TP and TN denote the number of correctly classified antigenically distinct and similar pairs, respectively. Conversely, FP and FN represent the number of misclassified as antigenically distinct and similar pairs, respectively. Furthermore, to robustly assess model performance on imbalanced datasets, we calculate the Area Under the Precision-Recall Curve (AUPRC_*k*_) for fold *k* or test year *k*, which examines Precision against Recall across varying classification thresholds. Here,

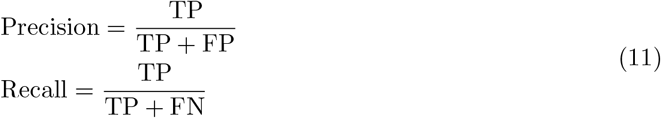

To obtain comprehensive performance estimates, the overall metrics are calculated as sample-size-weighted averages across all folds or test years:

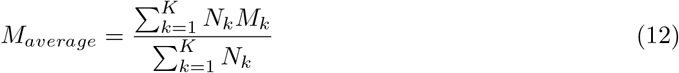

where *K* is the total number of evaluation partitions (folds or test years), *M*_*k*_ represents the metric value for partition *k*, and *N*_*k*_ denotes the number of antigen-antiserum pairs in the corresponding test set.

### Benchmarked models

We benchmark Vir3D against five established antigenic predictive models (MFPAD, AdaBoost, PREDAC, CNN-PSO, and PREDAC-CNN), three baselines (SVM, CNN, and GCN), and three sequence-encoding methods (AAindex, ESM-2, and ProtTrans). The implementation details, including the underlying principles and parameter configurations of these methods, are provided as follows:

#### MFPAD

MFPAD utilizes XGBoost to predict the antigenic distances of IAVs. It integrates binary-encoded HA1 sequences with four categories of features, which include the number of mutations, glycosylation sites, antigenic epitopes (A–E), and physicochemical properties. The hyperparameters for XGBoost are set as follows: `booster` = ‘gbtree’, `max_depth` = 13, and `n_estimators` = 200.

#### AdaBoost

AdaBoost utilizes an ensemble of decision trees for antigenic prediction by learning a nonlinear mapping from differences between pairwise HA1 sequences, which are encoded using amino acid mutation matrices from the AAindex2 database. The hyperparameters for AdaBoost are set as follows: `max_depth` = 1860, `max_features` = 0.394, and `learning_rate` = 1.393.

#### PREDAC

PREDAC utilizes a Naive Bayes classifier to discriminate antigenic variants from non-variants of IAVs. It integrates diverse features, including the differences between pairwise HA1 sequences in key antigenic regions (antigenic epitopes (A–E), receptor binding sites, and glycosylation sites), and physicochemical properties. The hyperparameters for the Naive Bayes classifier are set as follows: `alpha` = 1.0, `binarize` = 0.0, and `fit_prior` = True.

#### CNN-PSO

CNN-PSO utilizes a CNN optimized by Particle Swarm Optimization (PSO) to discriminate antigenic variants from non-variants. It utilizes a curated selection of amino acid substitution matrices (AAindex 2 and 3) to construct features from HA1 sequences. The core hyperparameters consist of three convolutional layers with `filters` = 193, 212, and 109, and `strides` = 3, 2, and 3.

#### PREDAC-CNN

PREDAC-CNN utilizes a one-dimensional CNN to discriminate antigenic variants from non-variants. It employs a ‘7-features’ encoding scheme for sequence representation, integrating six physicochemical properties from the AAindex1 database. The core hyperparameters are set as follows: `filters` = 6 and `kernel_size` = 3.

#### CNN

This baseline leverages a one-dimensional CNN to extract geometric features of the protein structure. Each residue is represented by a 184-dimensional feature vector, which is processed through two convolutional layers. These features are further enriched by incorporating learnable site-importance vectors, which are then condensed into a protein representation via a two-layer MLP operation. Subsequently, the protein representations of the virus pair are concatenated and passed through a transformer encoder layer for antigenic prediction. The core hyperparameters are set as follows: `kernel_size`=5, `stride` = 1, and `bias` = ‘True’.

#### SVM

This baseline employs an SVM for antigenic prediction of IAVs based on differences between pairwise HA1 sequences, which are encoded through amino acid mutation matrices from the AAindex2 database to capture physicochemical substitution effects. The hyperparameters for SVM are set as follows: `C` = 1.0, `kernel` = ‘rbf’, and `epsilon` = 0.1.

#### GCN

It utilizes a GCN to extract geometric embeddings from HA1 protein structures for antigenic prediction, where each residue is initially encoded into a 184-dimensional geometric embedding based on its coordinates. The model processes these geometric embeddings through a two-layer MLP operation, augments them with learnable site-importance embeddings, and then aggregates them through global mean pooling. The resulting protein representations are subsequently concatenated and fed into a transformer encoder layer and an MLP to predict antigenic distances.

#### AAindex

It maps each amino acid to a feature vector containing 566 distinct physicochemical properties from the AAindex1 database. To incorporate site-specific information, these feature vectors are augmented with learnable site-importance embeddings. The combined features are processed by a two-layer MLP to generate individual protein representations. Subsequently, the representations of paired viral proteins are concatenated and fed into a transformer encoder layer to explicitly model pairwise interactions, followed by an MLP for antigenic prediction.

#### ESM-2

It utilizes the esm2 t36 3B UR50D model to extract evolutionary features. It retrieves context-aware embeddings from the 36th hidden layer, yielding a 2560-dimensional representation for each site in the HA1 protein. These residue-level embeddings are further integrated with learnable site-importance embeddings and processed by a two-layer MLP to generate protein-level embeddings. Subsequently, the protein embeddings of two viruses are then concatenated and refined using a transformer encoder layer. Finally, an MLP is applied for antigenic prediction.

#### ProtTrans

It utilizes the ProtT5-XL-UniRef50 model to generate context-aware sequence embeddings. It extracts 1024-dimensional feature vectors from the model’s last hidden state, which are enhanced with learnable site-importance vectors and processed by a two-layer MLP to generate individual protein representations. The representations of paired viral proteins are subsequently concatenated and passed through a transformer encoder layer, followed by an MLP for antigenic prediction.

### Feature importance scores

By leveraging the attention mechanism inherent to the geometric graph learning framework, we calculate the importance score of each node by aggregating the attention weights of all edges connected to it. The resulting score quantifies the contribution of a given site in HA1 to viral antigenicity, with higher values indicating a more pronounced role in driving antigenic drift. Consequently, we prioritize the top 30 highest-scoring sites for downstream analysis.

### Statistical significance of region enrichment in top sites

To quantitatively evaluate the enrichment of Vir3D-prioritized sites within key antigenic regions known to affect the antigenic properties, we perform a hypergeometric test. Under the null hypothesis, the *P*-value represents the probability of observing at least *i* of the top 30 Vir3D-prioritized sites falling within a given antigenic region:

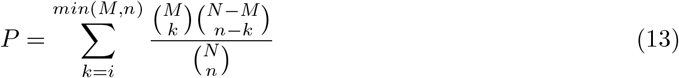

where *N* denotes the total length of the HA1 sequence (*N* = 328 for the H3 subtype and *N* = 317 for the H5 subtype), *M* is the total number of sites within the specific antigenic region, and *n* is the number of top-ranked important sites selected for analysis (*n* = 30 in this study). *P <* 0.05 indicates a statistically significant enrichment of model-identified key sites within the specific region.

### Structural visualization

To illustrate the spatial locations of the identified important sites, we map the top-ranked sites onto the corresponding HA1 structures and perform structural visualization using PyMOL (www.pymol.org) based on PDB entries 4KVN (A/Perth/16/2009) for H3 subtype and 4K62 (A/Indonesia/5/2005) for H5 subtype. To quantify the spatial proximity between the important sites and key antigenic regions known to affect the antigenic properties, we calculate the Euclidean distances between their *α*-carbon (C_*α*_) atoms. A distance threshold of 7 Å is applied. Specifically, an identified site is considered structurally associated with a key antigenic region known to affect the antigenic properties if the spatial distance between its C_*α*_ atom and that of any site within the key antigenic region is less than 7 Å.

### Antigenic map

To visualize antigenic drift of IAVs (Figure 3g, h, and Figure S3), we project IAVs of the H3 subtype from 2016–2018 and 2020–2022, as well as IAVs of the H5 subtype from 2013–2018 and 2015–2020, onto the antigenic maps using both FluNexus [**li2026flunexus**] https://flunexus.com/atmap.html and Racmacs [22]. FluNexus employs a manifold-based algorithm, while Racmacs utilizes multidimensional scaling to project HI titers into a two-dimensional space. To ensure reproducibility, the two-dimensional coordinates of antigens and antisera are obtained using over 1,000 optimizers with a fixed random seed of 0 for all maps.

### Definition of coverage rate

Influenza vaccine efficacy depends critically on the antigenic proximity between the selected vaccine strain and the viruses that will dominate the forthcoming epidemic. Because future epidemic strains are unknown at the time of selection yet characteristically emerge from the antigenic clusters circulating in the preceding season, the optimal selection strategy targets the antigenic centroid of the current virus landscape: the candidate strain that is antigenically similar to the greatest fraction of its contemporaries, and therefore most likely to confer broad cross-protective immunity against near-future variants. We operationalize this epidemiological principle through the coverage rate, defined for a candidate vaccine virus *v* over a reference set V_ref_ as:

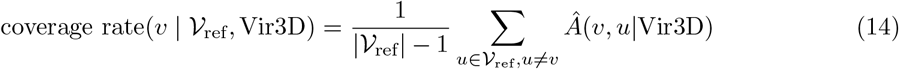

where *Â*(*v, u* | Vir3D) ∈ {0, 1} is a binary indicator of predicted antigenic similarity between candidate virus *v* and the reference virus *u*, as inferred by Vir3D. The coverage rate thus equals the fraction of reference viruses antigenically matched to *v*: a strain achieving a coverage rate approaching 1.0 is the antigenic centroid of V_ref_, and therefore the candidate best positioned to maximize population-level cross-protection.

Depending on the clinical objective, this general formulation is instantiated as two operationally distinct coverage rate types. For prospective vaccine selection, we define the predicted coverage rate: the reference set V_ref_ comprises candidate viruses collected in year *y* − 1, with year *y* denoting the target year, and Vir3D is trained exclusively on HI titer data available up to year *y* − 1. This strictly prospective design ensures that the metric reflects only information accessible at the time of an actual vaccine recommendation – no future data is exploited. For retrospective performance validation, we define the empirical coverage rate: the reference set V_ref_ consists of viruses that actually circulated during the target year *y*, and antigenic similarities are inferred by Vir3D trained on the complete dataset inclusive of year *y*. The empirical coverage rate thereby quantifies the true cross-protective efficacy of any recommended strain against the epidemic that actually materialized, and serves as the gold-standard benchmark for evaluating all three recommendation strategies (Vir3D, Oracle, and WHO).

To verify that the coverage rates derived from Vir3D-predicted antigenic distances faithfully reflect empirical coverage rates calculated from raw HI titers, we perform a comprehensive correlation analysis. To mitigate the statistical instability inherent in sparse single-year HI data, we aggregate antigen-antiserum pairs within a three-year temporal window, as WHO-recommended candidate vaccine strains for a target year are predominantly derived from viruses circulating within the preceding three years. The consistency between Vir3D-predicted coverage rates (*C*_*p,i*_) and HI-derived empirical coverage rates (*C*_*t,i*_) is quantified using the Pearson correlation coefficient *R* and the Spearman rank correlation coefficient *ρ*, defined respectively as:

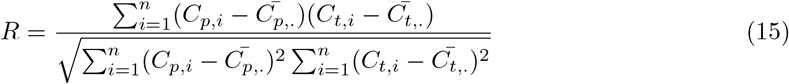

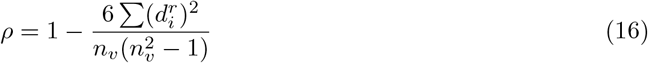

where 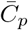 and 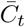 denote the sample means of the predicted and HI-derived coverage rates, respectively; 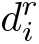 is the difference between the rank of *C*_*p,i*_ and the rank of *C*_*t,i*_ for the *i*-th virus; and *n*_*v*_ is the total number of virus pairs included in the analysis.

## Ethical Statements

The animal experimental protocols were reviewed and approved by the Committee on the Ethics of Animal Experiments at the Harbin Veterinary Research Institute, Chinese Academy of Agricultural Sciences (250220-04-GJ). All animal procedures were performed in accordance with the NIH Guide for the Care and Use of Laboratory Animals. Three-week-old male and female specific-pathogen-free (SPF) White Leghorn chickens were obtained from a certified commercial breeding supplier.

## Author Contributions

X.L. supervised the entire project, guided model design and experiments, and provided critical revisions to the manuscript. C.Z. developed the Vir3D framework, conducted all experiments, and wrote the manuscript. K.X. assisted with model validation and technical discussions. J.L., X.J., D.Z., L.C., Y.Li, and J.P. contributed to the literature review and provided valuable feedback during manuscript preparation. J.Z. assisted with model validation, provided valuable feedback during manuscript preparation, and provided critical revisions on the manuscript. Y.Liu provided supporting datasets, insightful suggestions on the experimental design, and result interpretation. X.S. provided overall academic support, critical revisions to the manuscript, and institutional resources. H.K. provided supporting datasets, critical revisions to the manuscript, and contributed to the experimental setup and model validation.

## Data Availability

The open-source Python implementation of Vir3D is available at https://github.com/xingyili/Vir3D. HA sequences of H3 influenza viruses can be retrieved from Smith et al. [22] and GISAID [36] at https://gisaid.org/. Corresponding HI data of H3 subtype can be downloaded at https://www.crick.ac.uk/research/platforms-and-facilities/worldwide-influenza-centre/annual-and-interim-reports. Additional HI data of H3 subtype can be compiled from the study of Smith et al. [22]. HA sequences of H5 influenza viruses can be retrieved from GISAID and GitHub at https://github.com/xingyili/FluNexus-methodbox/tree/main/H5_data. H5-specific HI data can be retrieved from GitHub at https://github.com/xingyili/FluNexus-methodbox/tree/main/H5_data. The HA1 structures of H3 (A/Perth/16/2009; PDB ID: [4KVN]) and H5 (A/Indonesia/5/2005; PDB ID: [4K62]) are downloaded from the Protein Data Bank (PDB) https://www.rcsb.org.

## Supplementary Materials

Supplementary Materials:

Figure S1. Framework of the five-fold cross-validation strategy. Figure S2. Framework of the retrospective testing strategy.

Figure S3. Antigenic cartography visualizing the antigenic drift using Racmacs. Figure S4. Validation of predicted coverage rates against HI-derived values.

Figure S5. Sequence logo shows the amino acid variation in the HA1 protein of U.S. dairy cow H5N1 viruses based on all available sequences deposited in GISAID (as of March 5, 2026).

Figure S6. Antigenic distances between representative H5 clade viruses and the TX/24 isolate, including its variants (TX/24-HA-88G and TX/24-HA-131M).

Figure S7. Overview of HI datasets for the H3 and H5 influenza A subtypes. Figure S8. Data distribution for the H3 subtype over influenza years (1968–2024). Figure S9. Data distribution for H5 subtype over influenza years (1996–2020).

Table S1. Annual coverage rates of vaccines recommended by WHO (2012–2024).

Table S2. Annual coverage rates of vaccines recommended by Vir3D and Oracle (2012–2024). Table S3. Representative H5 viruses from different clades included in this study and their corresponding specific isolates.

Table S4. HI titers obtained from cross-reactivity assays between representative H5 reference viruses and the TX/24 isolate and its variants.

