## Supplementary for "Structure-aware deep learning predicts influenza antigenicity and guides vaccine strain recommendation"

### 1. Supplementary Figures

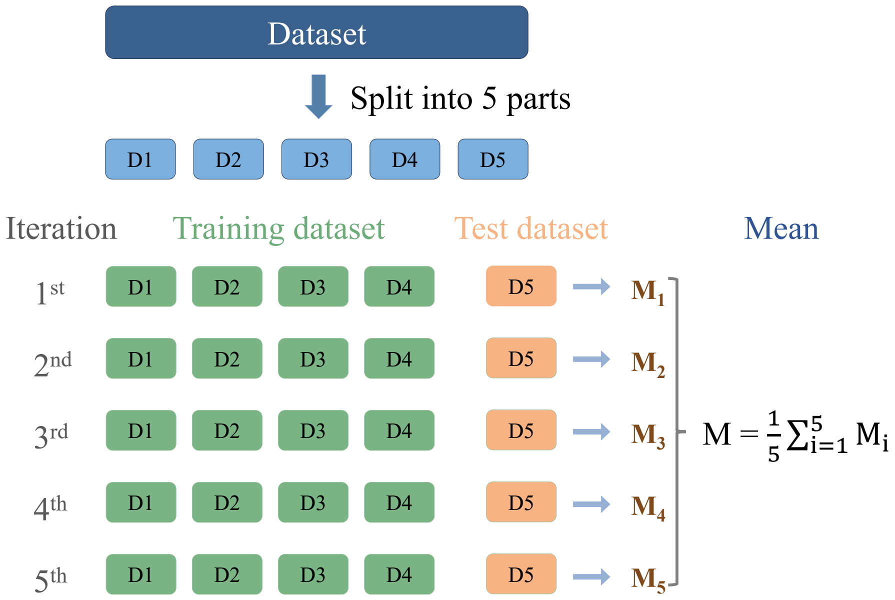


Figure S1. Framework of the five-fold cross-validation strategy. The complete dataset is randomly partitioned into five mutually exclusive subsets (D1–D5). Across five successive iterations, four subsets are combined to serve as the training dataset (green blocks) for model optimization, while the single remaining subset is held out as the test dataset (orange blocks) for performance evaluation. The overall model performance is determined by calculating the average ($M = \frac{1}{5}\sum_{i=1}^{5} M_i$) of the evaluation metrics ($M_i$) derived from all five independent iterations, providing a robust assessment of the model's predictive capability.

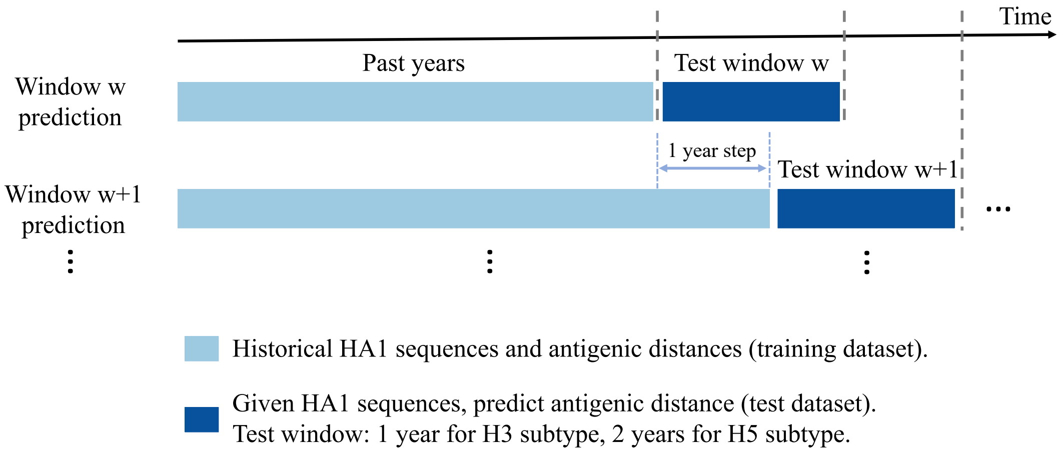


Figure S2. Framework of the retrospective testing strategy. For a given prediction window $w$, the model is trained on all accumulated historical HA1 sequences and their associated antigenic distances up to a specific time point (light blue bars). The trained model is subsequently evaluated on unseen viral data within a defined future test window (dark blue bars). To capture continuous viral evolution, the training–testing boundary advances iteratively with a fixed step size of 1 year. The duration of the test window is subtype-specific, spanning 1 year for the H3 subtype and 2 years for the H5 subtype.

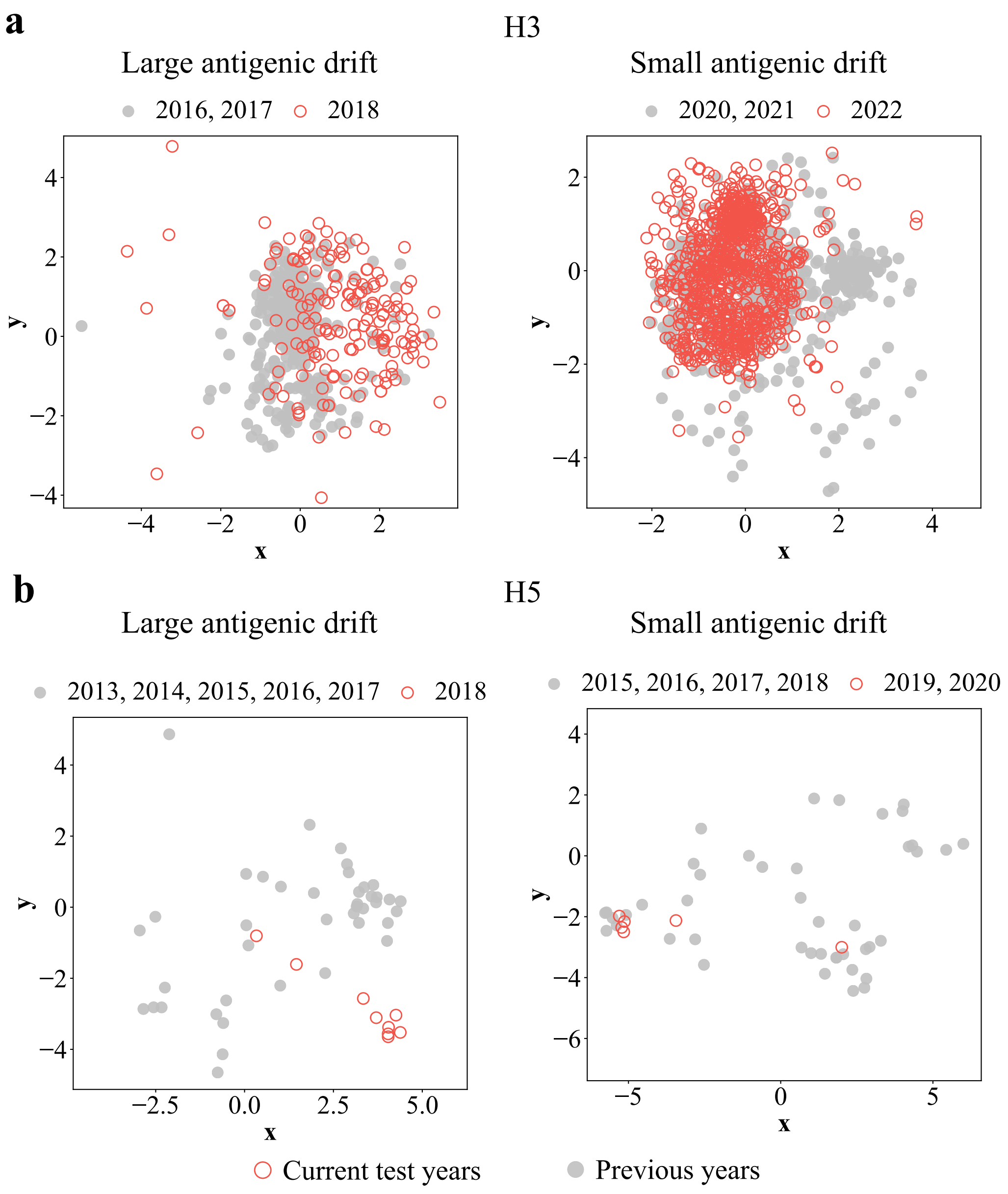


Figure S3. Antigenic cartography visualizing the antigenic drift using Racmacs. For each subtype, the map on the left shows an instance of large antigenic drift, while the map on the right shows an instance of small antigenic drift. (a) Antigenic cartography visualizing the antigenic drift of circulating viruses relative to those collected from the previous years for the H3 subtype. (b) Antigenic cartography visualizing the antigenic drift of circulating viruses relative to those collected from the previous years for the H5 subtype.

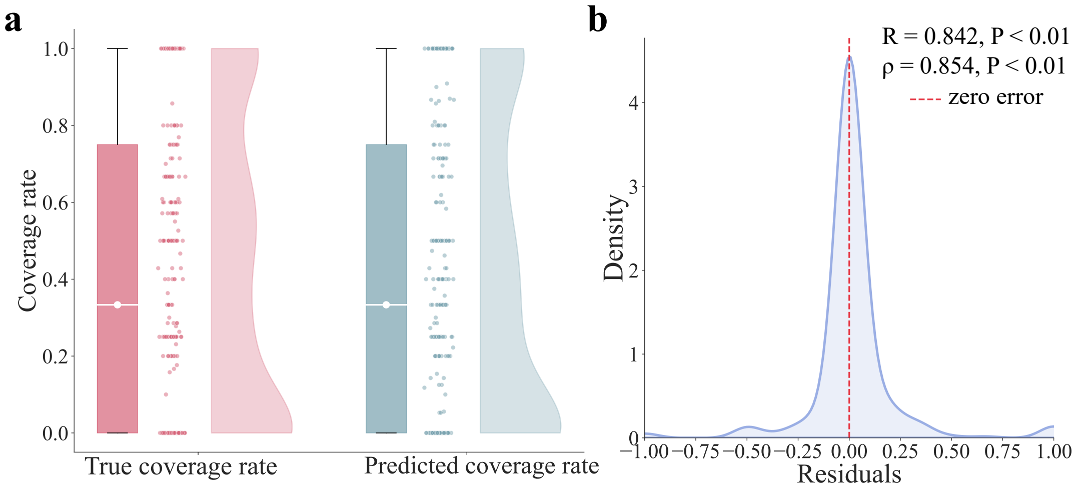


Figure S4. Validation of predicted coverage rates against HI-derived values. (a) Comparison of the distributions of coverage rates calculated based on HI titers and those inferred by Vir3D. The raincloud plots illustrate the consistency between the coverage rates derived from HI titers (pink, True coverage rate) and those predicted by Vir3D (blue, Predicted coverage rate). (b) Residual and correlation analysis. The density plot shows the distribution of residuals (True coverage rate - Predicted coverage rate), which is tightly centered around the zero-error dashed line. Statistical metrics indicate a strong positive correlation between predicted and observed values (Pearson $R = 0.842, P < 0.01$; Spearman $\rho = 0.854, P < 0.01$).

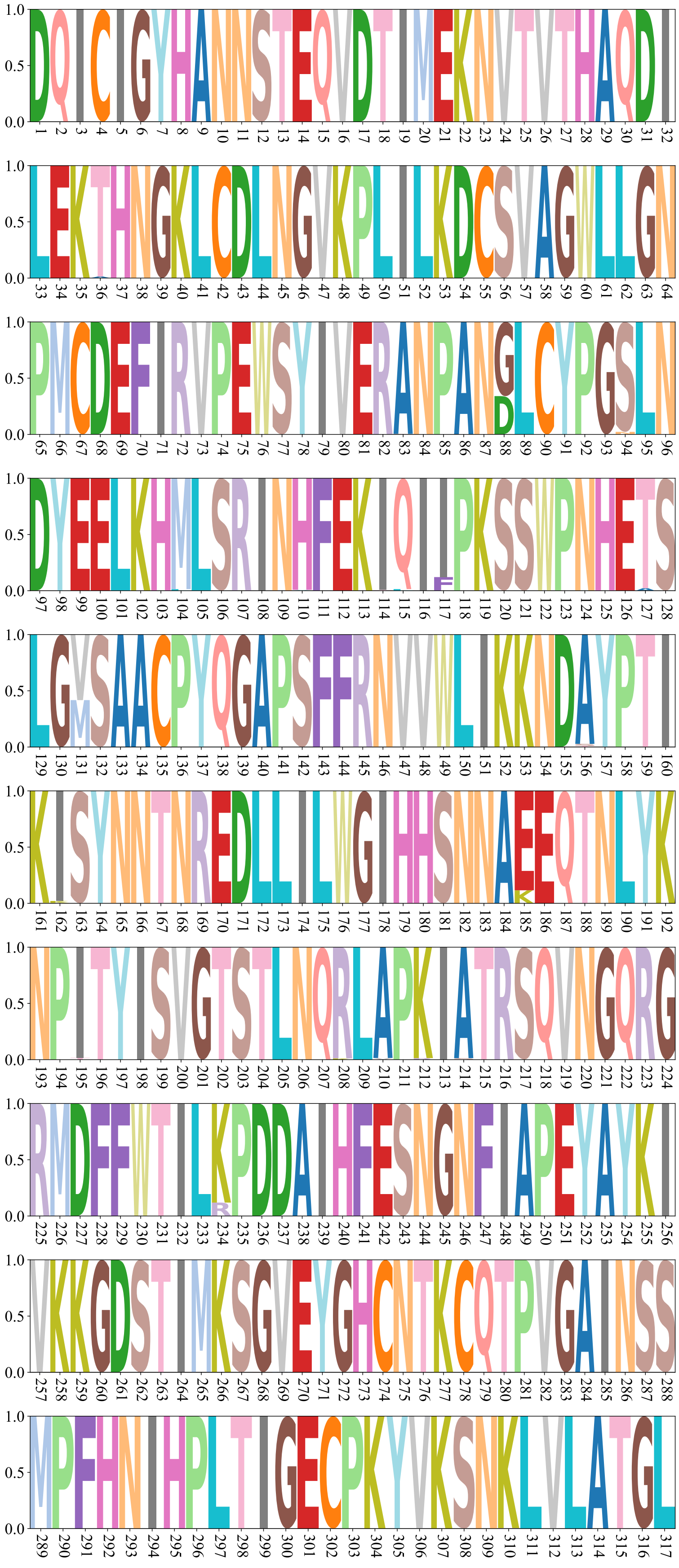

Figure S5. Sequence logo shows the amino acid variation in the HA1 protein of U.S. dairy cow H5N1 viruses based on all available sequences deposited in GISAID (as of March 5, 2026). The height of each letter represents the relative frequency of the corresponding amino acid at that position.

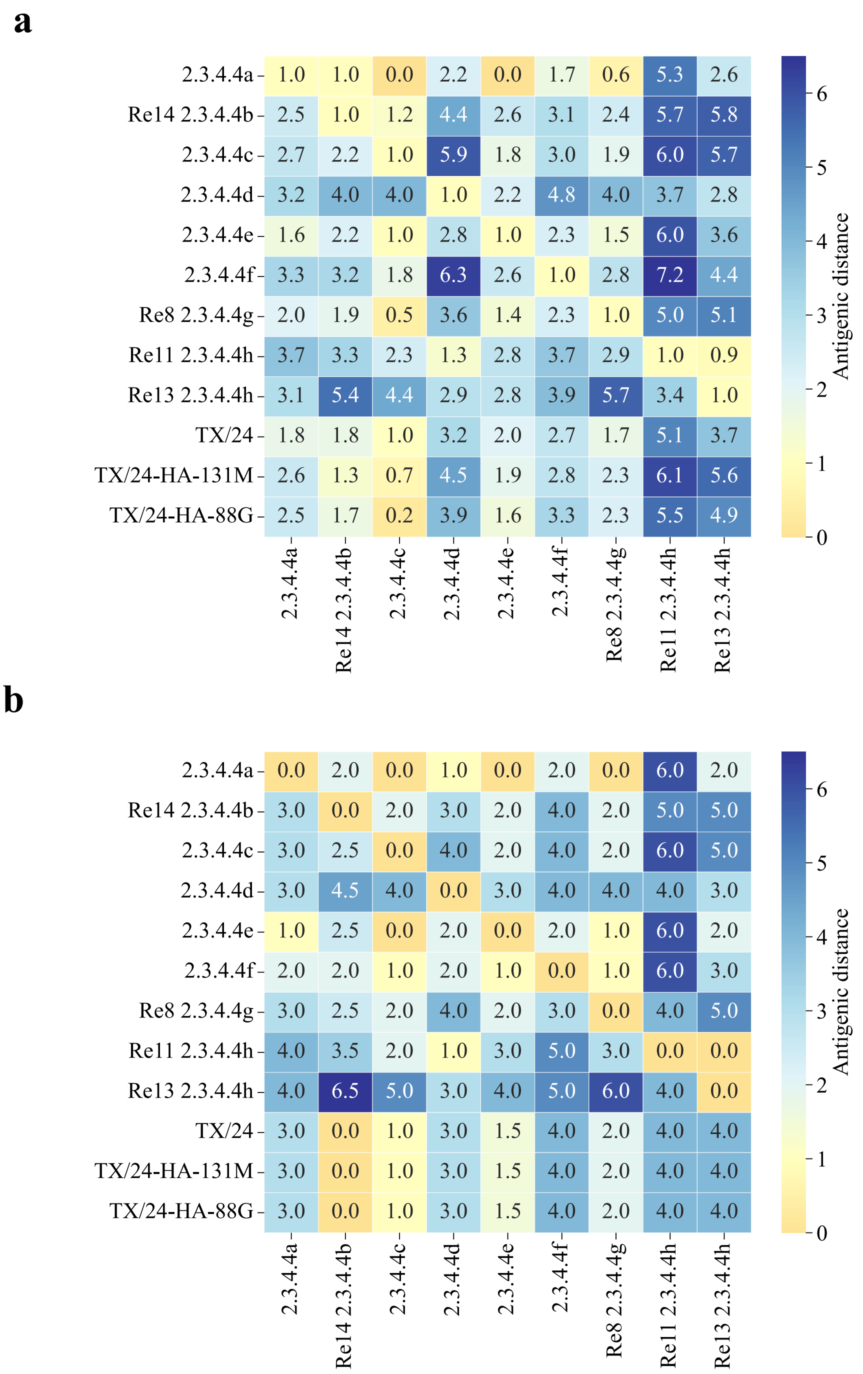


Figure S6. Antigenic distances between representative H5 clade viruses and the TX/24 isolate, including its variants (TX/24-HA-88G and TX/24-HA-131M). (a) Antigenic distances inferred by Vir3D. (b) Antigenic distances derived from HI assays.

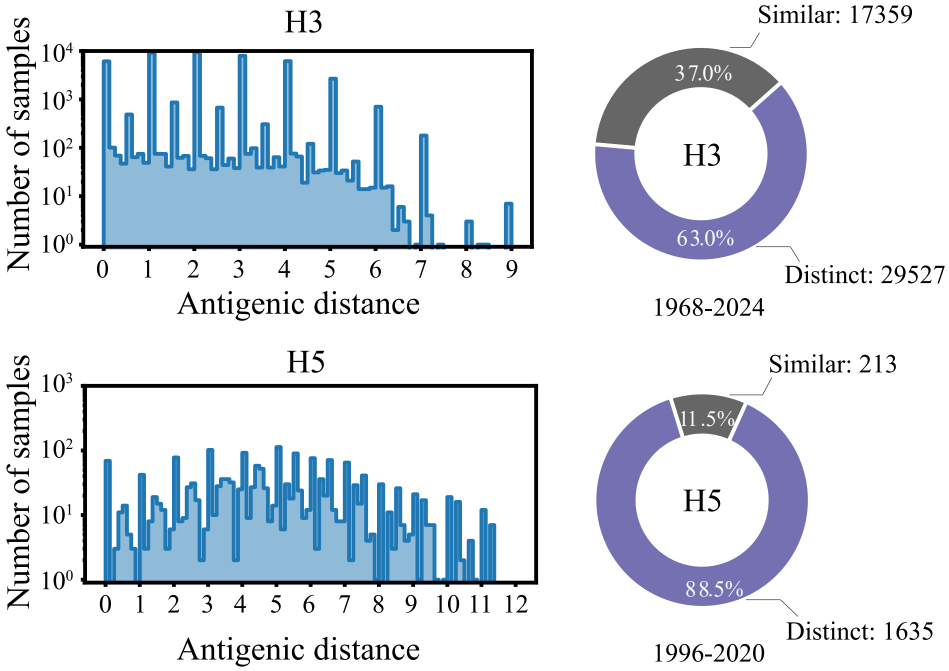


Figure S7. Overview of HI datasets for the H3 and H5 influenza A subtypes. (a) Distribution of the antigenic distances. (b) Distribution of the antigenically similar and distinct antigen-antiserum pairs.

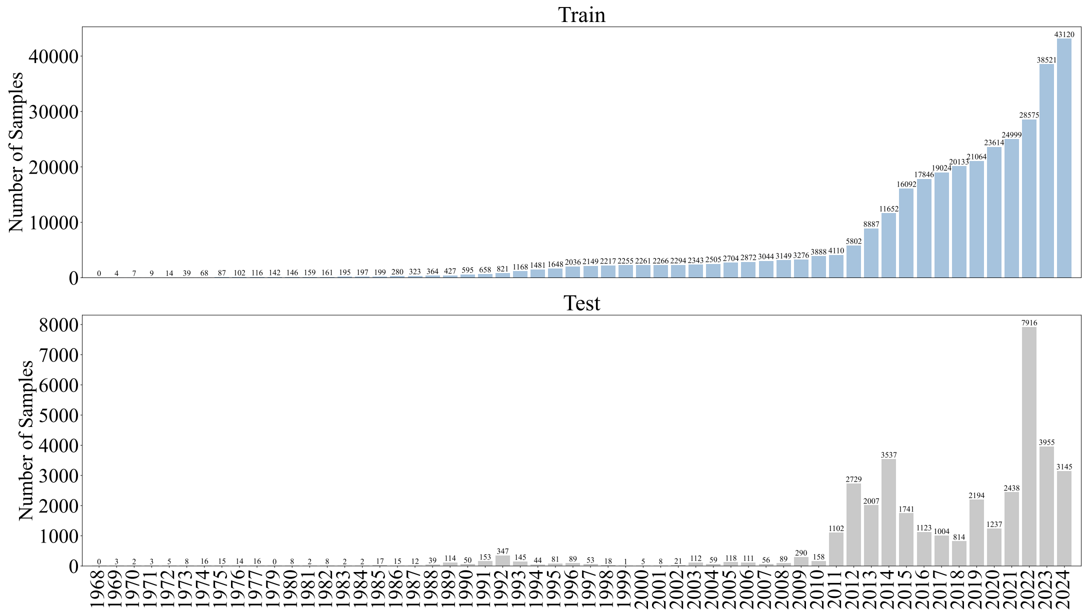


Figure S8. Data distribution for the H3 subtype over influenza years (1968–2024). The upper panel shows the number of antigen-antiserum pairs in the training dataset for each year, whereas the lower panel depicts the same for the test dataset.

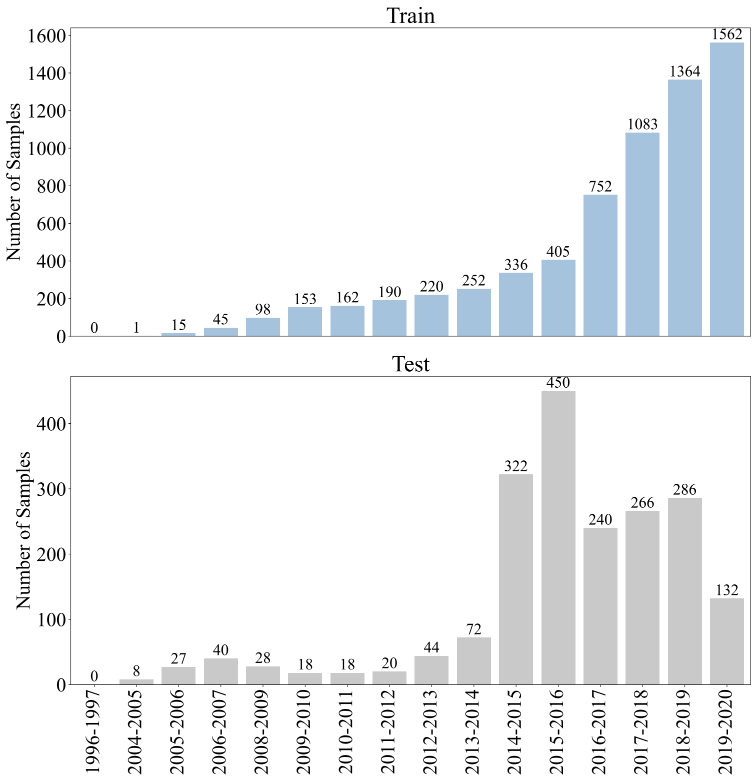


Figure S9. Data distribution for H5 subtype over influenza years (1996–2020). The upper panel shows the number of antigen-antiserum pairs in the training dataset for each year, whereas the lower panel depicts the same for the test dataset.

### 2. Supplementary Tables

Table S1. Annual coverage rates of vaccines recommended by WHO (2012–2024). (a) Coverage rates of vaccines recommended by WHO for the northern hemisphere. (b) Coverage rates of vaccines recommended by WHO for the southern hemisphere.

**a**

| Year | Vaccine | Coverage rate |
|---|---|---|
| 2024 | A/Thailand/8/2022\|EPI_ISL_16014504 | 0.97 |
| 2023 | A/Darwin/9/2021\|EPI_ISL_2233240 | 0.84 |
| 2022 | A/Darwin/9/2021\|EPI_ISL_2233240 | 0.80 |
| 2021 | A/Cambodia/e0826360/2020\|EPI_ISL_806547 | 0.98 |
| 2020 | A/Hong Kong/2671/2019\|EPI_ISL_20140890 | 0.75 |
| 2019 | A/Kansas/14/2017\|EPI_ISL_363642 | 0.88 |
| 2018 | A/Singapore/INFIMH-16-0019/2016\|EPI_ISL_291274 | 0.43 |
| 2017 | A/Hong Kong/4801/2014\|EPI_ISL_20140890 | 0.69 |
| 2016 | A/Hong Kong/4801/2014\|EPI_ISL_20140890 | 0.86 |
| 2015 | A/Switzerland/9715293/2013\|EPI_ISL_198223 | 0.88 |
| 2014 | A/Texas/50/2012\|EPI_ISL_129744 | 0.33 |
| 2013 | A/Texas/50/2012\|EPI_ISL_129744 | 0.94 |
| 2012 | A/Victoria/361/2011\|EPI_ISL_104004 | 0.95 |

**b**

| Year | Vaccine | Coverage rate |
|---|---|---|
| 2024 | A/Thailand/8/2022\|EPI_ISL_16014504 | 0.97 |
| 2023 | A/Darwin/9/2021\|EPI_ISL_2233240 | 0.84 |
| 2022 | A/Darwin/9/2021\|EPI_ISL_2233240 | 0.80 |
| 2021 | A/Hong Kong/2671/2019\|EPI_ISL_20140890 | 0.91 |
| 2020 | A/South Australia/34/2019\|EPI_ISL_20141035 | 0.85 |
| 2019 | A/Switzerland/8060/2017\|EPI_ISL_330887 | 0.74 |
| 2018 | A/Singapore/INFIMH-16-0019/2016\|EPI_ISL_291274 | 0.43 |
| 2017 | A/Hong Kong/4801/2014\|EPI_ISL_20140890 | 0.69 |
| 2016 | A/Hong Kong/4801/2014\|EPI_ISL_20140890 | 0.86 |
| 2015 | A/Switzerland/9715293/2013\|EPI_ISL_198223 | 0.88 |
| 2014 | A/Texas/50/2012\|EPI_ISL_129744 | 0.33 |
| 2013 | A/Victoria/361/2011\|EPI_ISL_104004 | 0.97 |
| 2012 | A/Perth/16/2009\|EPI_ISL_60761 | 0.78 |

Table S2. Annual coverage rates of vaccines recommended by Vir3D and Oracle (2012–2024). (a) Coverage rates of vaccines recommended by Vir3D. (b) Coverage rates of vaccines recommended by Oracle.

**a**

| Year | Vaccine | Coverage rate |
|---|---|---|
| 2024 | A/South Africa/R00721/2023\|Original | 0.99 |
| 2023 | A/Norway/20676/2022\|ClinicalSpecimen | 0.94 |
| 2022 | A/Nordrhein-Westfalen/1/2021\|Original | 0.88 |
| 2021 | A/Togo/827/2020\|Original | 0.97 |
| 2020 | A/Iceland/12621/2019\|SIAT1 | 0.86 |
| 2019 | A/England/80480634/2018\|original | 0.89 |
| 2018 | A/Greece/4/2017\|E10 | 0.70 |
| 2017 | A/RhodeIsland/30/2016\|Original | 0.75 |
| 2016 | A/SriLanka/61/2015\|E5/E1 | 0.99 |
| 2015 | A/SHIMANE/77/2014\|MDCK1+SIAT1 | 0.93 |
| 2014 | A/Boston/YGA_01169/2013\|P0 | 0.97 |
| 2013 | A/SYDNEY/195/2012\|MDCK2 | 0.94 |
| 2012 | A/Taiwan/1097/2011\|MDCK3+1 | 0.82 |

**b**

| Year | Vaccine | Coverage rate |
|---|---|---|
| 2024 | A/Kaliningrad/RII-1/2024\|MDCK-Siat1 | 0.99 |
| 2023 | A/South Africa/R00721/2023\|Original | 1.00 |
| 2022 | A/Norway/20676/2022\|ClinicalSpecimen | 0.91 |
| 2021 | A/Nordrhein-Westfalen/1/2021\|Original | 0.98 |
| 2020 | A/Togo/827/2020\|Original | 0.99 |
| 2019 | A/Iceland/12621/2019\|SIAT1 | 0.94 |
| 2018 | A/England/80480634/2018\|original | 1.00 |
| 2017 | A/Greece/4/2017\|E10 | 0.99 |
| 2016 | A/RhodeIsland/30/2016\|Original | 1.00 |
| 2015 | A/SriLanka/61/2015\|E5/E1 | 1.00 |
| 2014 | A/SHIMANE/77/2014\|MDCK1+SIAT1 | 0.99 |
| 2013 | A/Boston/YGA_01169/2013\|P0 | 0.99 |
| 2012 | A/SYDNEY/195/2012\|MDCK2 | 1.00 |

Table S3. Representative H5 viruses from different clades included in this study and their corresponding specific isolates.

| Virus | Specific isolate |
|---|---|
| 2.3.4.4a | DK/CQ/S1049/2014(H5N6) |
| Re14 2.3.4.4b | WS-SX-4-1-2020(H5N8) Re-14 |
| 2.3.4.4c | DK/ZJ/S1327/2016(H5N6) |
| 2.3.4.4d | CK/JX/5/2016(H5N6) |
| 2.3.4.4e | CK/HuN/6/2015(H5N6) |
| 2.3.4.4f | CK/HuN/1/2016(H5N6) |
| Re8 2.3.4.4g | A/chicken/Guizhou/4/2013(H5N1)Re-8 |
| Re11 2.3.4.4h | A/duck/Guizhou/S4184/2017(H5N6)Re-11 |
| Re13 2.3.4.4h | A/duck/Fujian/S1424/2020(H5N6)Re-13 |

Table S4. HI titers obtained from cross-reactivity assays between representative H5 reference viruses and the TX/24 isolate and its variants.

| Virus | 2.3.4.4a | Re14 2.3.4.4b | 2.3.4.4c | 2.3.4.4d | 2.3.4.4e | 2.3.4.4f | Re8 2.3.4.4g | Re11 2.3.4.4h | Re13 2.3.4.4h |
|---|---|---|---|---|---|---|---|---|---|
| 2.3.4.4a | 2048 | 91 | 512 | 512 | 2048 | 1024 | 2048 | 64 | 256 |
| Re14 2.3.4.4b | 256 | 362 | 128 | 128 | 512 | 256 | 512 | 128 | 32 |
| 2.3.4.4c | 256 | 64 | 512 | 64 | 512 | 256 | 512 | 64 | 32 |
| 2.3.4.4d | 256 | 16 | 32 | 1024 | 256 | 256 | 128 | 256 | 128 |
| 2.3.4.4e | 1024 | 64 | 512 | 256 | 2048 | 1024 | 1024 | 64 | 256 |
| 2.3.4.4f | 512 | 91 | 256 | 256 | 1024 | 4096 | 1024 | 64 | 128 |
| Re8 2.3.4.4g | 256 | 64 | 128 | 64 | 512 | 512 | 2048 | 256 | 32 |
| Re11 2.3.4.4h | 128 | 32 | 128 | 512 | 256 | 128 | 256 | 4096 | 1024 |
| Re13 2.3.4.4h | 128 | 4 | 16 | 128 | 128 | 128 | 32 | 256 | 1024 |
| TX/24 | 256 | 362 | 256 | 128 | 724 | 256 | 512 | 256 | 64 |
| TX/24-HA-131M | 256 | 362 | 256 | 128 | 724 | 256 | 512 | 256 | 64 |
| TX/24-HA-88G | 256 | 362 | 256 | 128 | 724 | 256 | 512 | 256 | 64 |
